# Outer membrane lipoprotein DolP interacts with the BAM complex and promotes fitness during envelope stress response

**DOI:** 10.1101/2020.09.23.308296

**Authors:** David Ranava, Yiying Yang, Luis Orenday-Tapia, François Rousset, Catherine Turlan, Lun Cui, Violette Morales, Cyril Moulin, Gladys Munoz, Jérôme Rech, Anne Caumont-Sarcos, Cécile Albenne, David Bikard, Raffaele Ieva

## Abstract

In Gram-negative bacteria, coordinated remodelling of the outer membrane (OM) and the peptidoglycan is crucial for envelope integrity. Envelope stress caused by unfolded OM proteins (OMPs) activates sigmaE (σ^E^) in Enterobacteria. σ^E^ upregulates OMP biogenesis factors, including the β-barrel assembly machinery (BAM) that catalyzes OMP-folding. Elevated σ^E^ activity, however, can be detrimental for OM integrity. Here we report that DolP (YraP), a σ^E^-upregulated OM lipoprotein important for envelope integrity, is a novel interactor of BAM and we demonstrate that OM-assembled BamA is a critical determinant of the BAM-DolP complex. Mid-cell recruitment of DolP had been previously associated to activation of septal peptidoglycan remodelling during cell division, but its role during envelope stress was unknown. We now show that DolP promotes cell fitness upon stress-induced activation of σ^E^ and opposes a detrimental effect caused by the overaccumulation of BAM in the OM. During envelope stress, DolP loses its association with the mid-cell, thus suggesting a possible link between envelope stress caused by impaired OMP biogenesis and the regulation of a late step of cell division.

## Introduction

The outer membrane (OM) of Gram-negative bacteria forms a protective barrier against harmful compounds, including several antimicrobials. This envelope structure surrounds the inner membrane and the periplasm that contains the peptidoglycan, a net-like structure made of glycan chains and interconnecting peptides. During cell division, the multi-layered envelope structure is remodelled by the divisome machinery ^1^. At a late step of division, septal peptidoglycan synthesized by the divisome undergoes splitting, initiating the formation of the new poles of adjacent daughter cells. Finally, remodelling of the OM barrier completes formation of the new poles in the cell offspring. The mechanisms by which cells coordinate OM remodelling with peptidoglycan splitting, preserving the permeability barrier of this protective membrane, are ill-defined ^2^.

Integral outer membrane proteins (OMPs) are crucial to maintain the OM permeability barrier. OMPs fold into amphipathic β-barrel structures that span the OM and carry out a variety of tasks. Porins are OMPs that facilitate the diffusion of small metabolites. Other OMPs function as cofactor transporters, secretory channels, or machineries for the assembly of proteins and lipopolysaccharide (LPS) ^3,4^, a structural component of the external OM leaflet that prevents the diffusion of noxious chemicals ^4^. The β-barrel assembly machinery (BAM) is a multi-subunit complex that mediates the folding and membrane insertion of OMPs transiting through the periplasm ^5,6^. The essential and evolutionarily conserved BamA insertase subunit is an OMP consisting of an amino (N)-terminal periplasmic domain made of polypeptide transport-associated (POTRA or P) motifs and a carboxy (C)-terminal 16-stranded β-barrel membrane domain that catalyzes OMP biogenesis ^7,8^. The flexible pairing of β-strands 1 and 16 of the BamA β-barrel controls a lateral gate connecting the interior of the barrel towards the surrounding lipid bilayer ^9–12^.

Conformational dynamics of the BamA β-barrel region proximal to the lateral gate is thought to locally increase the entropy of the surrounding lipid bilayer ^9,13,14^ and assist the insertion of nascent OMPs into the OM ^11,15,16^. The N-terminal periplasmic portion of BamA from the enterobacterium *Escherichia coli* contains five POTRA motifs that serve as a scaffold for four lipoproteins, BamBCDE, which assist BamA during OMP biogenesis ^17–19^. The N-terminal POTRA motif is also the docking site of the periplasmic chaperone SurA ^20^. Together with the chaperones Skp and DegP, SurA contributes to monitor unfolded OMPs transported into the periplasm by the inner membrane general secretory (Sec) pathway ^21,22^.

Defective OMP assembly causes periplasmic accumulation of unfolded protein transport intermediates. This envelope stress is signalled across the inner membrane to induce the sigmaE (σ^E^)-mediated transcriptional response ^23^. In the absence of stress, σ^E^ is sequestered by the inner membrane-spanning RseA factor. By-products of misfolded OMP turnover activate degradation of RseA, liberating σ^E 24^. The σ^E^ response copes with stress i) by upregulating genes involved in OMP biogenesis, such as the *bam* genes ^25^, and ii) by lowering the OMP biogenesis burden via a post-transcriptional mechanism ^26^. Whereas σ^E^ is essential ^27^, a tight control of cytosolic σ^E^ availability is necessary for optimal cell fitness and to prevent a potentially detrimental effect on the envelope ^28–31^. Remarkably, the functions of a number of genes upregulated by σ^E^ remains unknown. Among those, *yraP* encodes a ~20 kDa OM-anchored lipoprotein largely conserved in γ and β proteobacteria ^32–36^. YraP is crucial for OM integrity ^34^ and pathogenicity ^32^. A recent study showed that YraP localizes at the mid-cell during a late step of cell division, where it contributes to the regulation of septal peptidoglycan splitting by an unknown mechanism ^33^. These observations do not explain why YraP is upregulated in response to σ^E^ activation and how YraP helps coping with envelope stress.

During cell division, envelope stress caused by defective OMP biogenesis would place an additional burden on the already-complicated envelope reshaping process faced by cells. It remains unknown whether stress caused by unfolded OMPs influences envelope remodelling at the forming poles of two adjacent daughter cells. Here we present a functional investigation of *yraP* prompted by a genome-wide synthetic-defect screen. We demonstrate genetic interactions of *yraP* with *bam* genes and we provide evidence that YraP can associate with the BAM complex via an interaction with the BamA subunit. YraP is not critical for OMP assembly but promotes fitness upon activation of the σ^E^ response. Upon envelope stress or when BAM overaccumulates in the OM, YraP loses its association with the mid-cell, thus suggesting a possible link between the envelope stress response and septal peptidoglycan hydrolysis during a late step of cell division. Hence, we propose to rename YraP as DolP (<u>d</u>ivision- and <u>O</u>M stress-associated <u>l</u>ipoprotein).

## Results

### A genome-wide synthetic-defect screen identifies *dolP* genetic interactions

The mutant allele Δ*dolP::kan* ^37^ was introduced into *E. coli* BW25113 by P1 transduction. The resulting Δ*dolP* strain grew normally on LB medium (Figs. S1A and S1B), but was highly susceptible to vancomycin (Figs. 1A and S1B). This antibiotic is normally excluded from the OM of wild-type cells but inhibits growth of cells lacking OMP biogenesis factors such as *skp* and *surA* (Fig. S1B). The expression of C-terminally fused DolP protein variants in place of its wild-type form restored vancomycin resistance (Fig. 1A and Fig. S1C). This result supports the notion that DolP is important for envelope integrity ^33–36^. However, the role of DolP during envelope stress remains poorly understood.

**Fig. 1.**
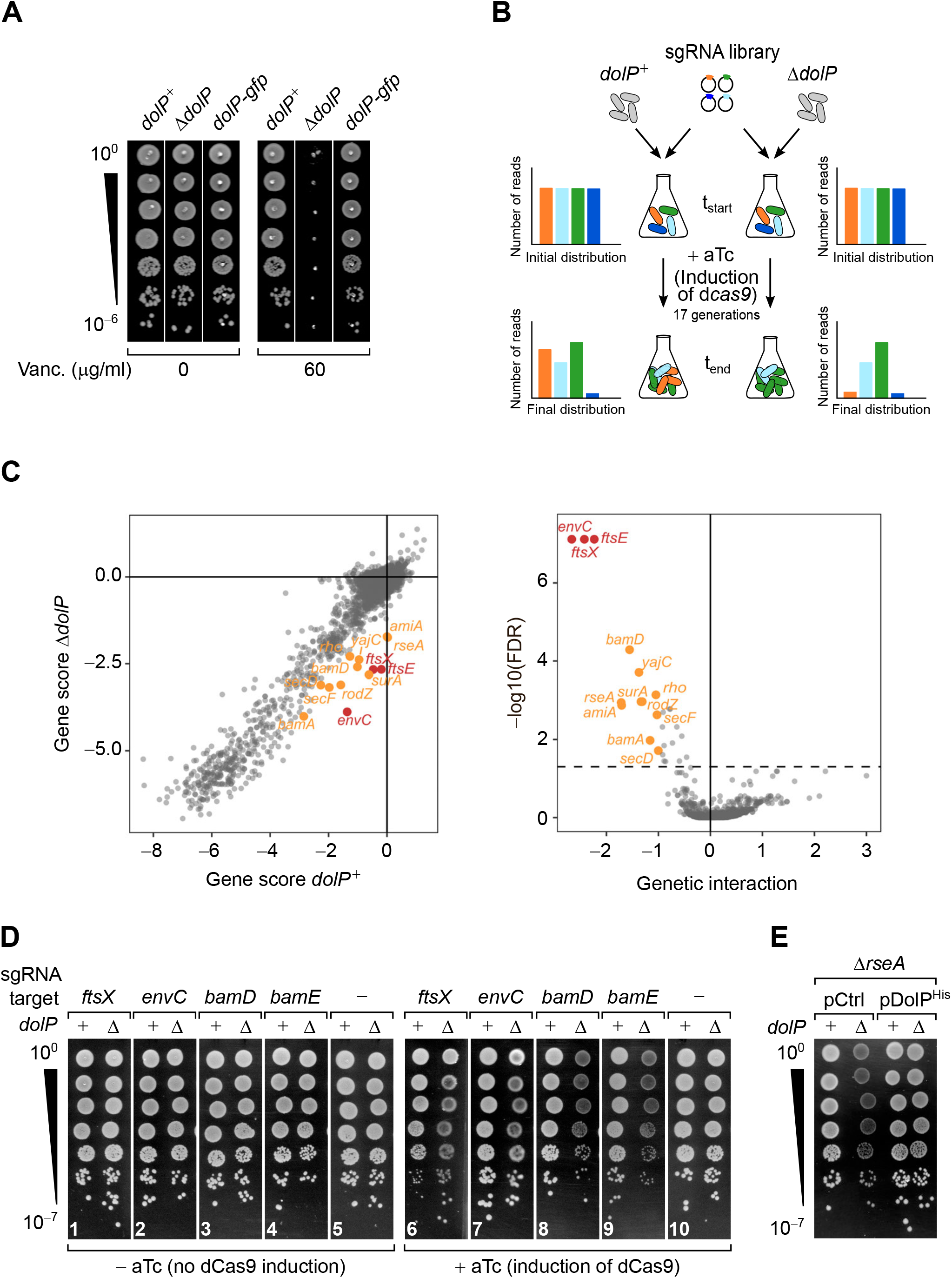
Genome-wide screen of *dolP* interactions. (**A**) The deletion of *dolP* impairs OM integrity. The indicated strains were serially diluted and spotted onto LB agar plates lacking or supplemented with 60 μg/ml vancomycin as indicated. (**B**) Schematic representation of the CRISPR-based gene silencing approach. LC-E75 (*dolP*^+^) or its Δ*dolP* derivative strain, both carrying d*cas9* under the control of an anhydrotetracycline (aTc)-inducible promoter in their chromosome were transformed with a library of plasmids encoding gene-specific sgRNAs. The library covers any *E. coli* MG1655 genetic features with an average of 5 sgRNAs per gene. Pooled transformed cells were cultured to early exponential phase prior to plasmid extraction and quantitative Illumina sequencing to assess the initial distribution of sgRNA constructs in each culture (t_start_). Upon addition of 1 μM aTc to induce sgRNA-mediated targeting of d*cas9* for approximately 17 generations, samples of cells from each culture were newly subjected to plasmid extraction and Illumina sequencing to determine the final distribution of sgRNA constructs (t_end_). (**C**) Left: Comparison of gene scores obtained in *dolP^+^* and Δ*dolP* screens. The log2 fold-change (log2FC) between t_end_ and t_start_ calculated for each sgRNAs (Fig. S2B) was grouped by gene target, and their median (Table S1) was used to derive fitness gene scores. Right: Volcano plot of the *dolP* genetic interaction (GI) scores. The x-axis shows a genetic interaction score calculated for each gene based on the minimum hypergeometric (mHG) test conducted on the ranked difference of sgRNA-specific log2FC values between the Δ*dolP* and the *dolP*^+^ screens. The y-axis shows the log10 of the false discovery rate (FDR) of the test. The dashed line shows FDR = 0.05. In both panels, genes highlighted in orange have FDR < 0.05 and GI>1 whereas genes highlighted in red have FDR < 0.05 and GI>2. (**D** and **E**) Validation of the genetic interactions determined in (C). (D) LC-E75 (*dolP*^+^) or its Δ*dolP* derivative strain expressing sgRNAs that target the indicated genes were serially diluted and spotted on LB agar lacking or supplemented with aTc to induce expression of d*cas9*, as indicated. (E) BW25113 derivative cells deleted of *rseA* or both *rseA* and *dolP* were transformed with an empty vector (pCtrl) or a plasmid encoding DolP (pDolP^His^). Ectopic expression of DolP^His^ was driven by the leaky transcriptional activity of P_*trc*_ in the absence of IPTG. (D and E) 10-fold serial dilutions of the indicated transformants were spotted on LB agar.

A previous study showed that lack of DolP causes a severe growth defect in a *surA* deletion strain, however the cause of this synthetic phenotype has remained unclear ^34^. To gain insights into the role of DolP, we subjected Δ*dolP* cells to a genome-wide synthetic-defect screen exploiting a Clustered Regularly Interspersed Short Palindromic Repeat interference (CRISPRi) approach. Targeting of the catalytically inactive dCas9 nuclease by gene-specific single guides RNAs (sgRNAs) enables gene repression (Fig. 1B) ^38^. The EcoWG1 sgRNA library targeting the entire genome of *E. coli* MG1655 ^39^ was introduced into isogenic Δ*dolP* or *dolP*^+^ MG1655-derivative strains. The fitness of each knockdown was then compared in these backgrounds by deep-sequencing of the sgRNA library after ~17 growth generations. The outputs obtained from two independent tests were highly reproducible (Fig. S2A). A strong fitness defect in the Δ*dolP* strain was caused by the targeting of *envC* (Fig. 1C, Fig. S2B and Tables S1 and S2), followed by the targeting of *ftsX* and *ftsE* (Fig. 1C, Tables S1 and S2). A validation growth test showed that the synthetic fitness defect observed for Δ*dolP* cells was caused by dCas9-dependent silencing of *ftsX* and *envC* (Fig. 1D, panels 6 and 7). The ABC transporter-like complex FtsE/FtsX has multiple roles in organizing the cell divisome, including the recruitment of periplasmic EnvC, a LytM domain-containing factor required for the activation of amidases that hydrolyse septal peptidoglycan ^40^. This peptidoglycan remodelling reaction is mediated by two sets of highly controlled and partially redundant amidases, AmiA/AmiB and AmiC ^41,42^. Whereas AmiA and AmiB are activated at the inner membrane/peptidoglycan interface by the coordinated action of FtsE/FtsX and EnvC, activation of AmiC requires the OM-anchored LytM domain-containing lipoprotein NlpD ^43,44^. Under laboratory conditions, the activity of only one of these two pathways is sufficient for septal peptidoglycan splitting, whereas inhibition of both pathways leads to formation of chains of partially divided cells, *i.e.* cells that have begun to divide but are blocked at the step of septal peptidoglycan splitting ^44^. A recent report showed that *dolP* is necessary to complete septal peptidoglycan splitting and to promote cell separation when the AmiA/AmiB pathway is inactive, somehow linking DolP to AmiC activation ^33^. Thus, the reduced fitness caused by silencing of *envC, ftsE or ftsX* in Δ*dolP* cells (Fig. 1C) may be explained by impaired cell separation when both the AmiA/AmiB and the AmiC pathways are not active. In keeping with this notion, *amiA* itself was found among the negative fitness hits of the CRISPRi screen (Fig. 1C). *amiB* was not a hit (Table S2), probably because AmiA is sufficient to split septal peptidoglycan in the absence of other amidases ^45^.

Most importantly, the CRISPRi approach identified novel *dolP*-genetic interactions that had a score similar to that obtained for *amiA* (Fig. 1C). These included an interaction with *rseA*, encoding the inner membrane σ^E^-sequestering factor, as well as with *bamD*, encoding an essential subunit of the BAM complex ^34,46^. In accordance with the screen output, a serial dilution assay confirmed that silencing of *bamD* (as well as *bamE*, encoding a stoichiometric interactor of BamD) causes a fitness defect in cells lacking DolP (Fig. 1D, panels 8 and 9). In addition, the interaction of *dolP* with *rseA* was confirmed in the genetic background of a BW25113 strain (Fig. 1E). Further genes, involved in OMP biogenesis and more generally in protein secretion, had a lower interaction score (Table S2 and Fig. S2C) and are highlighted in Fig. 1C. These comprise *bamA*, the OMP chaperone-encoding gene *surA*, as well as the genes encoding the Sec ancillary complex SecDF-YajC that contributes to efficient secretion of proteins including OMPs ^22^ and that was shown to interact with the BAM complex ^47,48^. Collectively, the results of the CRISPRi screen indicate that the function of DolP is particularly critical for cell fitness upon inactivation of septal peptidoglycan hydrolysis by AmiA, as well as when the assembly of proteins in the OM is impaired.

### DolP improves cell fitness when the OM undergoes stress

At a first glance, the newly identified genetic interaction between *dolP* and *bamD* (Figs. 1C and 1D) points to a possible role of DolP in OMP biogenesis. However, the overall protein profile of the crude envelope fraction was not affected by the deletion of *dolP* (Fig. S3A, lanes 1-4). OMPs, such as the very abundant OmpA and OmpC ^49^, can be recognized by their characteristic heat-modifiable migration patterns when separated by SDS-PAGE ^50^ (Fig. S3 A-D). The levels of OmpA (Fig. S3B), LamB (Fig. S3C) and OmpC (Fig. S3D) were not affected in Δ*dolP* samples. Similarly, the assembly kinetics of an autotransporter OMP was not impaired (Fig. S3E). Importantly, the levels of proteins encoded by σ^E^-upregulated genes, such as BamA and BamE, were not increased in Δ*dolP* cells (Fig. 2A), indicating a wild-type-like σ^E^ activity in this strain. The envelope protein profiles were not affected also when *dolP* was deleted in cells lacking one of the OMP periplasmic chaperones Skp or DegP (Fig. S3A, lanes 5-12), suggesting that DolP does not play a function redundant with that of these periplasmic factors. Taken together, these observations suggest that DolP is not crucial for efficient OMP biogenesis. The genetic interaction of *dolP* with *bamD* and *surA* (Fig. 1C) may also suggest that DolP is required for optimal survival when the OM undergoes modifications caused by activation of the σ^E^ response ^34,46,51^. This hypothesis is supported by the observation that σ^E^ activation caused by *rseA* silencing or deletion worsens the fitness of Δ*dolP* cells (Figs. 1C and 1E).

**Fig. 2.**
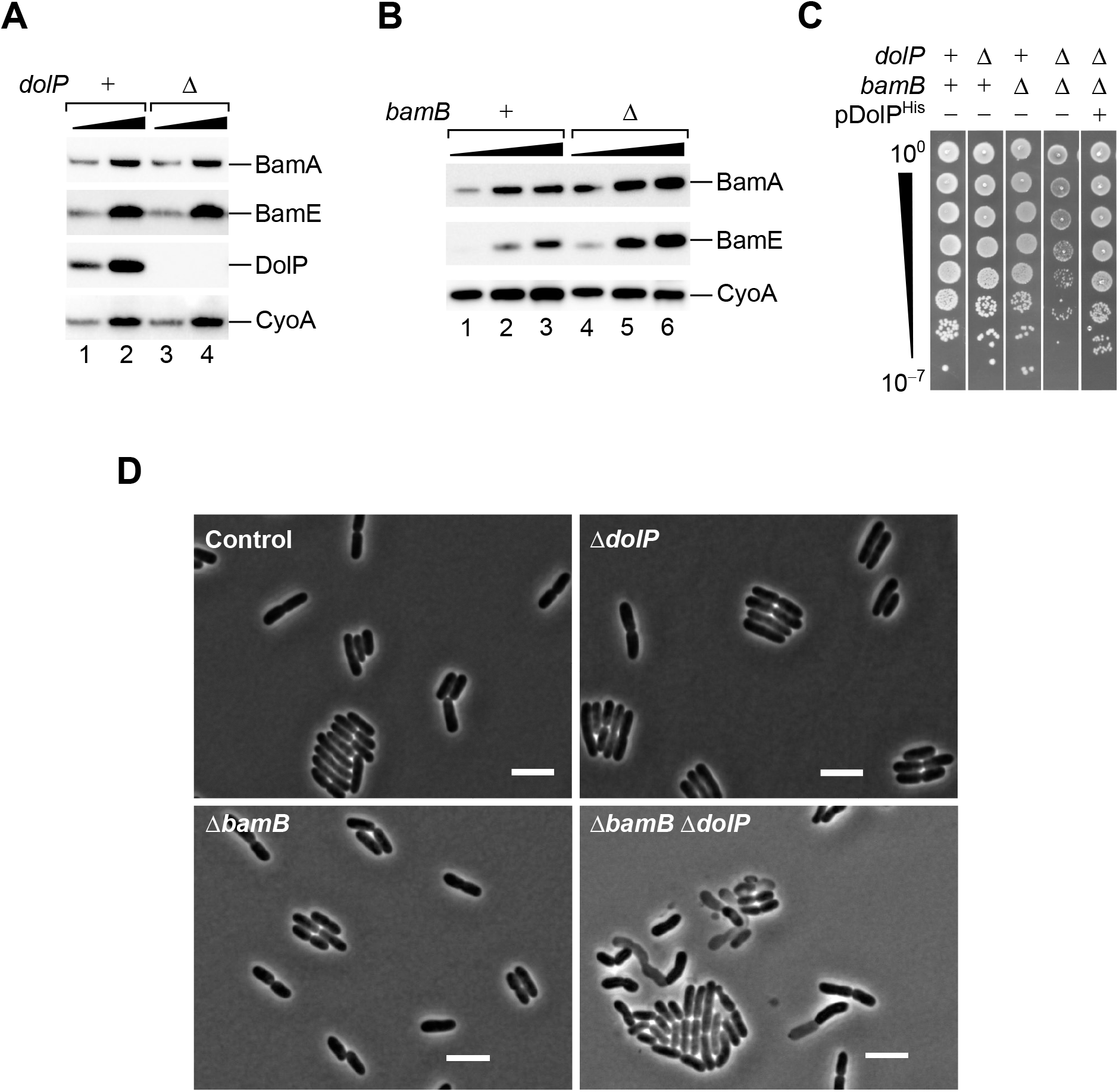
DolP promotes fitness in cells that undergo envelope stress. (**A**) One and three folds of a sample containing the total cell lysate fraction from a BW25113 (*dolP^+^*) strain and a derivative Δ*dolP* strain were analyzed by SDS-PAGE and immunoblotting using the indicated antisera. (**B**) One, two and three folds of a sample containing the total cell lysate fraction from a BW25113 (*bamB^+^*) strain and a derivative Δ*bamB* strain were analyzed by SDS-PAGE and immunoblotting using the indicated antisera. (**C**) BW25113 and derivative cells deleted of *dolP, bamB* or both genes were cultured, serially diluted and spotted on LB agar. A BW25113 derivative strain deleted of both *dolP* and *bamB* and transformed with pDolP^His^ was cultured, serially diluted and spotted on LB agar supplemented with ampicillin. (**D**) Overnight cultures of BW25113 (control), Δ*dolP*, Δ*bamB and* Δ*dolP* Δ*bamB*, were freshly diluted in LB medium and re-incubated at 30°C until OD_600_ = 0.3. Cell were visualized on 1% (w/v) agarose pads by phase contrast microscopy. Bar = 5 μm.

To further test the importance of DolP under envelope stress conditions, we deleted *dolP* in a strain lacking *bamB*, which was identified with a lower genetic score by the CRISPRi approach (Table S2). In Δ*bamB* cells, the σ^E^ response is partially activated ^19,52^, causing the upregulation of *bam* genes (Fig. 2B). A strain carrying the simultaneous deletion of *dolP* and *bamB* was viable but growth-defective. Normal growth was restored by ectopic expression of a C-terminally polyhistidine tagged DolP protein variant (Fig. 2C). The Δ*bamB* envelope protein profile presented a marked reduction of the major heat-modifiable OMPs, OmpA and OmpC, however the concomitant lack of DolP did not enhance this OMP defect (Fig. S3A, lanes 13-18). Notably, phase-contrast analysis of the same Δ*dolP* Δ*bamB* strain revealed a number of cells with altered morphology (Fig. 2D). This result suggests that the reduced fitness of Δ*dolP* Δ*bamB* cells cannot be ascribed to an exacerbation of the OMP biogenesis defect already caused by lack of BamB, pointing to a different role of DolP under envelope stress conditions.

### DolP promotes growth in cells with increased levels of BAM complex

The σ^E^ response leads to alteration of the OM protein content, with an increase of the level of the BAM complex and a reduction of numerous other proteins, including the most abundant OMPs and some lipoproteins ^25,53,54^. To gain further insights into the role of DolP during activation of the σ^E^ response, we explored the effect of prolonged overproduction of BAM. To this end, the genes encoding wild-type BamABCD and a C-terminally polyhistidine-tagged BamE protein variant were ectopically expressed via the isopropylthiogalactoside (IPTG)-inducible *trc* promoter (P_*trc*_) as a transcriptional unit, adapting a previously established method ^55^. With 400 μM IPTG, the concentration of BAM complex that accumulated in the membrane fraction was roughly similar to the concentration of the major OMPs OmpA or OmpC (Fig. 3A, lane 2). Importantly, we noticed that prolonged BAM overproduction caused a partial detrimental effect in the wild-type BW25113 strain (Fig. 3B). The excess of BamA was responsible for impaired growth, as the excess of different subsets of BAM subunits that did not include BamA or an excess of OmpA obtained using a similar overproduction plasmid (see also the subsequent description of Fig. S7B) had no detectable effects in our growth tests (Fig. 3B). The detrimental effect of BAM overproduction was caused by the overaccumulation of BamA in the OM, as the overproduction of an assembly-defective BamA variant, BamA^ΔP1^, that lacks the N-terminal POTRA1 motif and largely accumulates in the periplasm ^20^, did not impair growth to the same extent (Figs. S4A-C). Even a slight increment of BAM, due to the leaky transcriptional activity of the *trc* promoter (Fig. 3A, lane 1) caused a mild but noticeable deterioration of the OM permeability barrier to vancomycin (Fig. 3C), suggesting that a small increment in BAM levels can enhance the permeability of the OM. Strikingly, in the absence of vancomycin, we noticed that the detrimental effect caused by IPTG-induced OM overaccumulation of BAM was more severe in a Δ*dolP* strain (Fig. 3D). This difference was particularly evident with 200 μM IPTG, which had only a minor inhibitory effect on the growth of wild-type cells but strongly impaired the growth of a Δ*dolP* strain. Similar to *dolP*, *skp* is upregulated by σ^E^ and its deletion causes sensitivity to vancomycin (Fig. S1B). In contrast to Δ*dolP*, a Δ*skp* strain harbouring the same BAM overproduction plasmid could grow as the wild-type reference (Fig. 3D), suggesting a specific effect of DolP in supporting growth when BAM is overproduced.

**Fig. 3.**
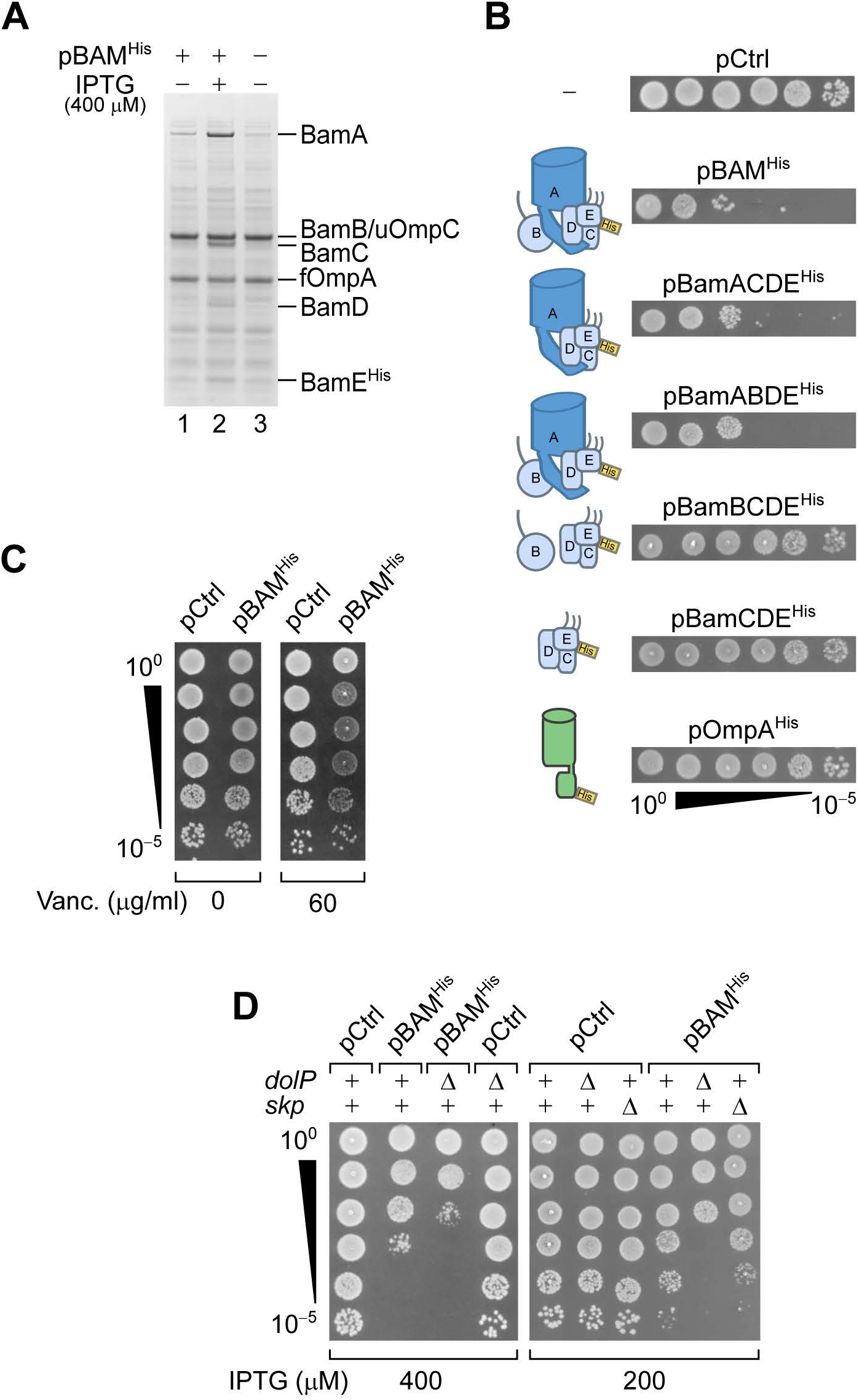
DolP opposes an envelope detrimental effect caused by BAM overaccumulation. (**A**) BW25113 cells harbouring pBAM^His^ as indicated were cultured and supplemented with no IPTG or 400 μM IPTG for 1 hour prior to collecting cells. The protein contents of the envelope fractions were analysed by SDS-PAGE and coomassie staining. Prior to loading, samples were heated for 5 min at 90°C, a temperature which is not sufficient to fully denature OmpA (folded OmpA, fOmpA). The band of BamB overlaps with the band of the major porin unfolded OmpC (uOmpC). (**B**) BW25113 cells carrying a control empty vector (pCtrl), or the indicated plasmids for ectopic overproduction of BAM or subsets of BAM subunits or OmpA were serially diluted and spotted on LB agar containing 400 μM IPTG. The diagrams show the overproduces proteins. (**C**) Wild-type BW25113 cells carrying an empty control vector (pCtrl) or pBAM^His^ were serially diluted and spotted on LB agar lacking or supplemented with 60 μg/ml vancomycin. No IPTG was added to these medium. (**D**) The BW25113 and the derivative Δ*dolP* or Δ*skp* strains carrying an empty control vector (pCtrl) or pBAM^His^ were serially diluted and spotted on LB agar supplemented with 400 or 200 μM IPTG, as indicated.

### DolP interacts with BamA assembled in the OM

The observed interaction between *dolP* and *bam* genes prompted us to investigate whether DolP physically interacts with the BAM complex. To this end, a construct encoding C-terminally protein A-tagged DolP was ectopically expressed in Δ*dolP* cells. The envelope of cells expressing DolP^ProtA^ was solubilized using digitonin as main mild-detergent component prior to IgG-affinity chromatography (Fig. 4A, coomassie staining). Site-specific enzymatic cleavage of an amino acid linker between DolP and the protein A tag was used for protein elution. Notably, BamA, BamC, BamD and BamE were immunodetected in the elution fraction of protein A-tagged DolP (Fig. 4A, lane 3). In contrast, the inner membrane protein CyoA and cytosolic RpoB were not detected. Next, BAM^ProtA^ (consisting of wild-type BamABCD and a C-terminally protein A-tagged BamE protein variant) was ectopically overproduced to isolate the BAM complex via IgG-affinity purification (Fig. 4B, Coomassie staining). Importantly, in addition to the BamE bait and other subunits of the BAM complex, DolP was also immunodetected in the elution fraction (Fig. 4B, lane 3). Other proteins of the bacterial envelope (Skp, OmpA, and F_1_β of the F_1_F_0_ ATP synthase) or cytosolic RpoB were not detected (Fig. 4B). Taken together, our native pull-down analysis indicates that DolP and BAM have affinity for each other.

**Fig. 4.**
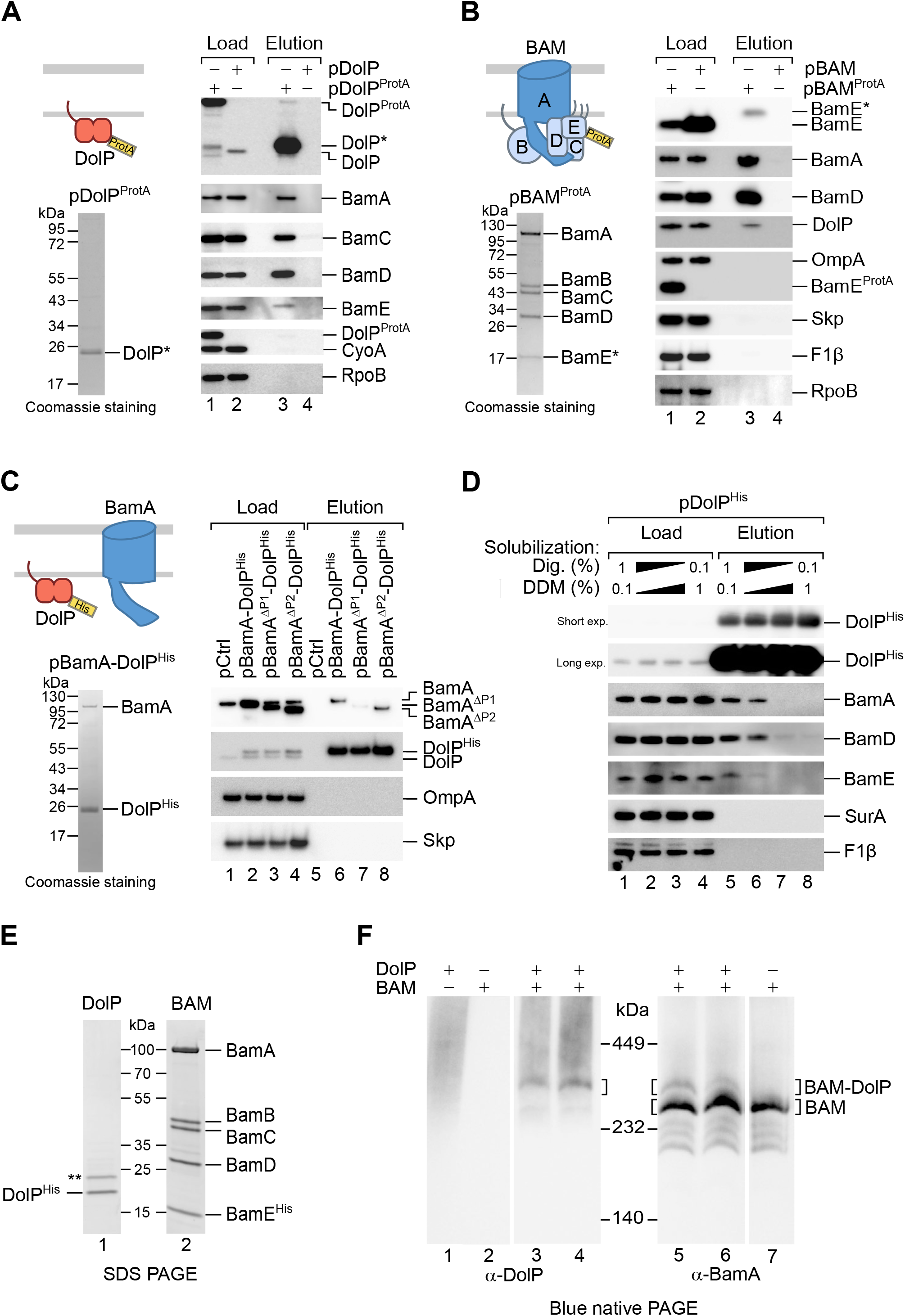
DolP associates with the BAM complex via an interaction with BamA. (**A**, **B**). The envelope fractions of BW25113 cells carrying the indicated plasmids were solubilized with 1% (w/v) digitonin and 0.1% (w/v) DDM and subjected to IgG affinity purification of either protein A-tagged DolP (A) or protein A-tagged BamE (B). The load and elution fractions were analyzed by SDS-PAGE. The coomassie staining of the elution of protein A-tagged DolP (A) or protein A-tagged BamE (B) are shown below the diagrams representing overproduced proteins. Blotted protein from load and elution fractions were detected by immunolabeling using the indicated antisera. Load 0.5% (A) 1% (B); Elution 100%. The asterisk indicates the products of TEV digestion obtained with samples containing DolP^ProtA^ (A) or BamE^ProtA^ (B). (**C**) The envelope fractions of BW25113 cells carrying the plasmids overproducing His-tagged DolP and the indicated BamA protein variants (deleted of POTRA1 or of POTRA2) were solubilized with 1% (w/v) digitonin and 0.1% (w/v) DDM and subjected to Ni-affinity purification. The load and elution fractions were analyzed by SDS-PAGE. The coomassie staining of the elution of His-tagged DolP overproduced together with wild-type BamA is shown below the diagram representing overproduced proteins. Blotted protein from load and elution fractions were detected by immunolabeling using the indicated antisera. Load 2%; Elution 100%. (**D**) Equal aliquots of the envelope fraction of BW25113 cells expressing His-tagged DolP were solubilized with the indicated concentrations (w/v) of digitonin and DDM: 1%, 0.1% (lane 1), 0.8%, 0.3% (lane 2), 0.3%, 0.8% (lane 3), 0.1%, 1% (lane 4). His-tagged DolP was purified by Ni-affinity chromatography. In all cases, proteins were eluted in the presence of 0.3% (w/v) digitonin and 0.03% (w/v) DDM. Load 0.2%; Elution 100%. (**E**) The envelope fraction of BW25113 cells overproducing DolP^His^ or the BAM complex containing C-terminally His-tagged BamE was subjected to protein extraction with 1% (w/v) DDM and Ni-affinity purification and gel filtration chromatographies. The elution fractions were analyzed by SDS-PAGE and coomassie staining. The double asterisk indicates a contaminant protein in the elution of DolP. (**F**) Roughly equimolar quantities of purified His-tagged BAM complex and DolP were incubated alone for 1 hour at 4°C (lanes 1, 2 and 7), or together for 1 hour at 4°C (lanes 3 and 6) or for 30 min at 25°C (lanes 4 and 5), prior to blue native-PAGE and immunoblotting using the indicated antisera.

To explore whether the central BAM subunit, BamA, is a critical determinant of the BAM-DolP interaction, we performed Ni-affinity purification using the solubilized envelope fraction obtained from cells overproducing BamA and C-terminally polyhistidine-tagged DolP. Under these conditions, BamA was efficiently co-eluted together with DolP^His^, demonstrating that BamA and DolP can interact even in the absence of stoichiometric amounts of the BAM lipoproteins (Fig. 4C, coomassie staining). To assess if the interaction of DolP and BAM takes place at the OM, DolP^His^ was overproduced together with the assembly-defective form BamA^ΔP1^. When expressed together with DolP^His^, assembly-defective BamA^ΔP1^ was highly depleted in the corresponding eluate (Fig. 4C, lane 7), even though BamA^ΔP1^ was only marginally reduced in the crude envelope fraction with respect to wild-type BamA (Fig. 4C, lane 3). In contrast to BamA^ΔP1^, the BamA^ΔP2^ variant, which is efficiently assembled into the OM (Fig. S4), was co-eluted to a similar extent as wild-type BamA (Fig. 4C, lane 8). This result indicates that DolP has affinity for OM-assembled BamA.

In seeking a detergent that would interfere with the interaction of BAM and DolP, and allow their purification as separate components, we solubilized the envelope fraction with increasing amounts of n-dodecyl β-D-maltoside (DDM), a detergent previously used to isolate the native BAM complex ^55^. At concentrations of DDM between 0.3% (w/v) and 1% (w/v), we observed a drastic reduction in the amounts of BAM subunits that were co-eluted with DolP^His^ (Fig. 4D), indicating that the BAM-DolP interaction is sensitive to DDM. We thus used 1% (w/v) DDM to extract and purify His-tagged DolP or His-tagged BAM as separate components (Fig. 4E). When analyzed by blue native-PAGE and immunoblotting, purified DolP gave rise to a diffused signal at around 450 kDa (Fig. 4F, lane 1), suggesting a dynamic multimeric organization of this protein. Purified BAM migrated as expected at 250 kDa (Fig. 4F, lane 7). When roughly equimolar amounts of purified BAM and DolP were pre-incubated in the presence of a low DDM concentration and subsequently resolved by blue native-PAGE, a complex with an apparent molecular weight higher than that of the BAM complex was detected with both anti-BamA and anti-DolP specific antibodies (Fig. 4F, lanes 3 to 6). Taken together our results demonstrate that the BAM complex and DolP interact to form a larger complex and that OM-assembled BamA is a critical determinant of this interaction. Only a portion of BAM and DolP associate to form the BAM-DolP complex, indicating that the interaction is substoichiometric.

### BamA overaccumulation in the OM reduces DolP mid-cell localization

In light of our observation that DolP interacts with BAM, we asked whether the envelope localization patterns of DolP and BAM are reciprocally linked. First, we monitored the effect of DolP expression on the localization of the chromosomally encoded BamD^mCherry^ subunit of the BAM complex. This protein generated a fluorescence signal throughout the envelope that was not affected by the lack or the overproduction of DolP (Fig. S5). Next, we checked the effect of BAM overaccumulation in the OM on DolP localization. DolP associates with the OM and accumulates at mid-cell during a late step of cell division ^33^. To monitor the localization of DolP, we used a strain harbouring a chromosomal *dolP-gfp* fusion (Fig. 1A). The localization of the DolP^GFP^ fusion protein (Fig. S6A) was analyzed concomitantly with other two chromosomally-encoded markers of the division septum, ZipA^mCherry^ or NlpD^mCherry^. ZipA is involved in an early step of divisome assembly and accumulates at division sites before, as well as, during envelope constriction (Fig. S6B) ^56^. Instead, NlpD is a late marker of cell division involved in the activation of AmiC and accumulates at septa that are already undergoing constriction (Fig. S6C) ^42,44^. DolP^GFP^ accumulated at mid-cell sites where the envelope appeared invaginated, showing a localization pattern similar to that of NlpD^mCherry^ (Fig. S6C) ^33^. We investigated the effect of short-lived (1 hour) BAM overproduction on DolP^GFP^ localization. Strikingly, we found that BAM overproduction depleted DolP^GFP^ from mid-cell sites (Figs. 5A, center, and S8C). In contrast, no obvious effects on cell division nor on mid-cell recruitment of ZipA^mCherry^ and NlpD^mCherry^ was observed (Fig. S7A). The overproduction of BamA alone was sufficient to alter the distribution of the DolP^GFP^ fluorescence signal in constricting cells, reducing its intensity at constriction sites and enhancing it at decentred positions along the cell axis (Fig. 5B, right plot). In contrast, the overproduction of only the four BAM lipoproteins (Fig. 5A, right) as well as the overproduction of OmpA (Fig. S7B) had no obvious effects on DolP mid-cell localization.

**Fig. 5.**
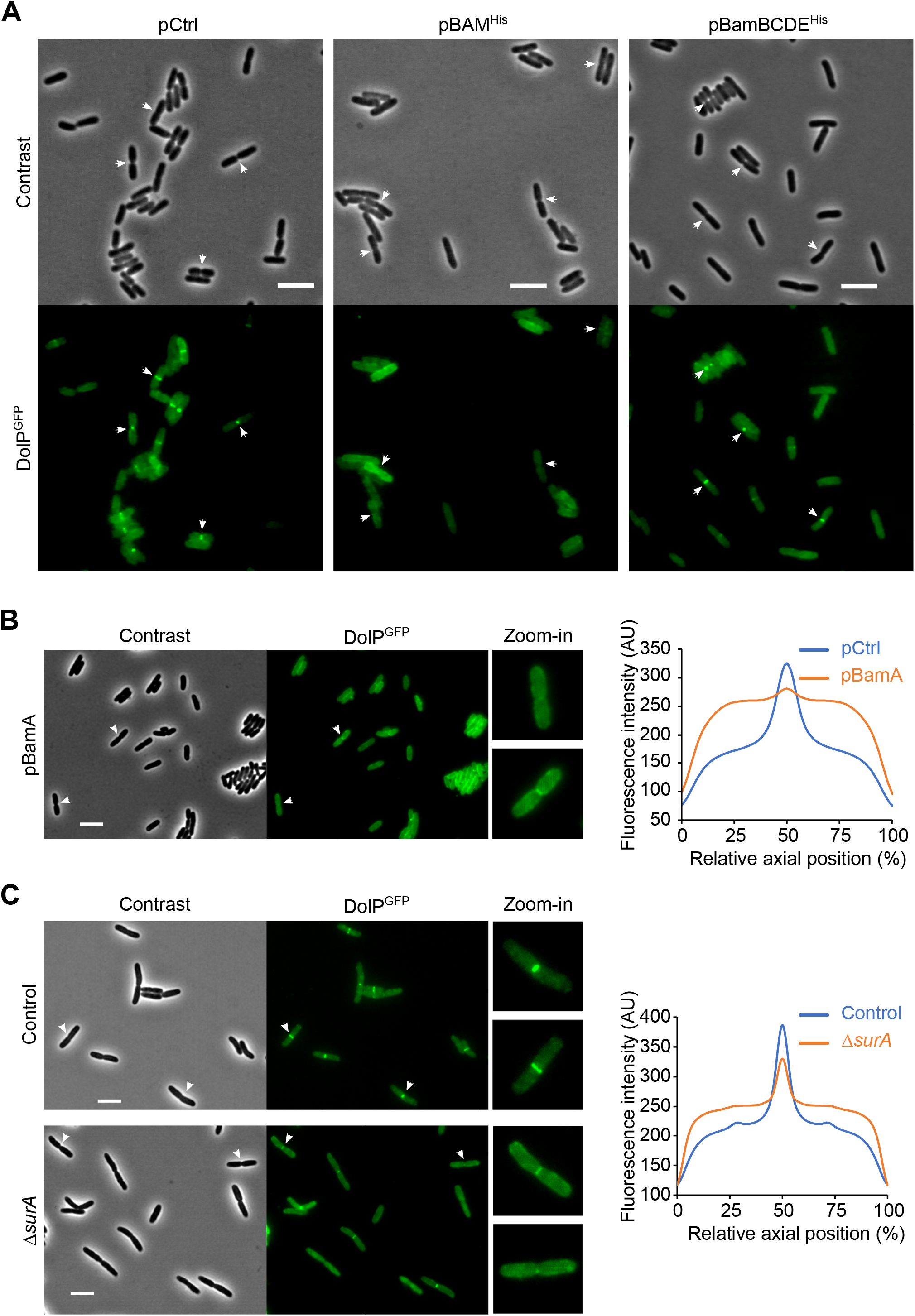
BamA overaccumulation in the OM and envelope stress interfere with mid-cell localization of DolP. (**A**) Overnight cultures of BW25113 cells harboring the chromosomal fusion *dolP-gfp* and transformed with either pCtrl (empty vector) or pBAM^His^, or pBamBCDE^His^ were freshly diluted in minimal M9 medium, incubated at 30°C until OD_600_ = 0.1 and supplemented with 400 μM IPTG for 1 hour. Cell samples were visualized on 1% (w/v) agarose pads by phase contrast and fluorescence microscopy. Arrowheads indicate envelope constriction sites between forming daughter cells. Bar = 5 μm. (**B**) Left: Overnight cultures of BW25113 cells harboring the chromosomal fusion *dolP-gfp* and transformed with either pCtrl (empty vector) or pBamA were cultured and visualized as in (A). Bar = 5 μm. Right: The collective profiles of fluorescence distribution versus the relative position along the cell axis were plotted for cells transformed with pBamA (orange) and cells transformed with pCtrl (blue). Only cells with a constriction (187, pBamA; 361, pCtrl) were taken into account for the collective profile plots. (**C**) Left: Overnight cultures of BW25113 (control) or Δ*surA* derivative cells carrying the *dolP-gfp* chromosomal fusion were freshly diluted in LB medium and incubated at 30°C until OD_600_ = 0.3. Cell samples were visualized as in (A). Bar = 5 μm. Right: The collective profiles of fluorescence distribution versus the relative position along the cell axis is shown for Δ*surA* cells (orange) and *surA*^+^ control cells (blue). Only cells with a constriction (320, Δ*surA*; 318, Control) were taken into account for the collective profile plots.

As BAM catalyzes OMP assembly, we asked whether this activity interferes with DolP mid-cell localization. To address this question, we made use of an inactive BamA mutant form (Fig. S8A) harbouring a polyhistidine peptide extension at its C-terminal β-strand ^57^. Similar to the overaccumulation of the BAM complex or BamA, the overaccumulation of BamA^His^ interfered with DolP mid-cell localization (Figs. S8B and S8C), without affecting ZipA^mCherry^ and NlpD^mCherry^ (Fig. S7C), indicating that the cellular localization of DolP does not depend on the OMP-assembly activity of BamA. In contrast, the ability of BamA to assemble into the OM was a critical determinant of the observed septal depletion of DolP. In fact, the periplasm-accumulating BamA^ΔP1/His^ variant (Fig. S8D) did not impair DolP^GFP^ mid-cell localization (Fig. S8B, center), whereas the OM-overaccumulating BamA^ΔP2/His^ did (Figs. S8B, right, S8C, and S8D). Taken together, these results suggest that the overaccumulation of BamA in the OM interferes with the recruitment of DolP at mid-cell sites.

### DolP mid-cell localization is impaired under envelope stress conditions

As DolP is critical for fitness under envelope stress conditions, we asked whether the localization of DolP^GFP^ would be affected in mutants undergoing envelope stress. To this end, we analyzed the localization of DolP^GFP^ in strains lacking either the OMP chaperone SurA or the lipoprotein BamB. Both Δ*surA* and Δ*bamB* strains are defective in OMP biogenesis and produce higher levels of BAM complex due to activation of the σ^E^ response ^19,51,52,58^. Importantly, the frequency of mid-cell labelling by DolP^GFP^ was reduced in Δ*surA* cells both in minimal (Fig. S9A) and LB (Fig. 5C) culture media. In contrast, lack of SurA did not affect septal recruitment of the late cell division marker NlpD (Fig. S9B). The analysis of the fluorescence plot profiles of constricted cells clearly showed a marked reduction of the DolP^GFP^ signal at mid-cell sites and higher fluorescence levels in decentred positions along the cell axis (Fig. 5C, right plot). As for the Δ*surA* strain, DolP^GFP^ accumulated at the mid-cell with a lower frequency when *bamB* was deleted (Fig. S9C). Together, these results indicate that the depletion of DolP at mid-cell sites occurs during envelope stress-induced activation of the σ^E^ response.

## Discussion

Despite its critical role in maintaining OM integrity, the reason why DolP is upregulated by σ^E^ has remained unclear ^32–34^. Here we demonstrate that DolP interacts with the BAM complex and is crucial for fitness in cells with elevated σ^E^ activity and when BAM overaccumulates in the OM. Our results have important implications for the understanding of the mechanisms that coordinate OM and peptidoglycan remodelling and help preserving envelope integrity.

A previous study ^34^ has shown that the concomitant deletion of *dolP* and *surA* generates a severe synthetic growth defect. However, the role of DolP during envelope stress and in OMP biogenesis had remained unclear ^34,36^. Here we demonstrate that DolP interacts with the BamA subunits of the BAM complex. DolP, however, is not a stoichiometric component of the BAM complex and thus this interaction is probably transient or has a lower affinity compared to the interactions between BamA and the lipoproteins of the BAM complex. Several pieces of evidence presented in this study suggest that DolP is not strictly required for OMP assembly into the OM. In addition, we show that the deletion of *dolP* does not cause envelope stress and activation of the σ^E^ response. Increased σ^E^ activity has been reported to alleviate growth defects caused by the absence or inactivation of OMP biogenesis factors ^59–62^. On the contrary, activation of σ^E^ caused a growth defect in a Δ*dolP* strain. This result suggests that, under envelope stress conditions, DolP plays a fitness role that is not directly related to OMP biogenesis. More specifically, we have obtained evidence that DolP opposes a detrimental effect that can derive from the overaccumulation of BAM complex in the OM.

Further studies are warranted to determine the molecular bases of how DolP promotes fitness during activation of the σ^E^ response. It is worth noting, however, that DolP consists of two BON (bacterial OsmY and nodulation) domains, folding motifs that have been proposed to interact with hydrophobic ligands such as lipids ^63^. Phospholipid-binding by DolP has been reported in a recent study ^64^. A key aspect of our results hints at a possible role of the lipid-binding function of DolP in improving the fitness of cells that undergo envelope stress. We have shown that DolP interacts with OM-assembled BamA, a subunit of the BAM complex that is predicted to interfere with the organization of the surrounding lipid bilayer and to generate an energetically favourable environment for the insertion of nascent OMPs in the OM ^65,66^. BamA-mediated membrane destabilization was shown by molecular dynamics simulations, as well as by reconstituting BamA into proteoliposomes ^9,67^. Here, we have provided evidence that the progressive overaccumulation of BAM in the OM has an increasingly detrimental effect, enhancing the OM permeability at low increments and impairing growth at higher levels. Our observation that DolP is needed for optimal growth when the BAM complex overaccumulates in the OM is compatible with a role of DolP in membrane stabilization at BAM sites. It remains to be seen how DolP docks to BamA and whether it influences the organization of BAM sites and of the surrounding lipid bilayer.

Another key finding of our study is that DolP septal localization is sensitive to envelope stress conditions. During envelope stress, the OM undergoes a significant alteration of its protein composition. Activation of the σ^E^ response triggers activation of genes encoding OMP biogenesis factors, but also the posttranscriptional downregulation of many OMPs including porins and OmpA ^25^. OMPs are largely arranged in clusters ^68–70^ embedded by highly organized LPS molecules in the external leaflet of the OM ^4^ and the rigidity of their β-barrel structures contributes to the mechanical stiffness of the OM ^71^. Posttranscriptional downregulation targets of σ^E^ activation also include the lipoproteins Pal and Lpp ^53,54^, which are critical for OM integrity ^72,73^. Whereas we have shown that BAM overaccumulation in the OM influences DolP localization, it is possible that other modifications of the OM during envelope stress contribute to impair the recruitment of DolP at constriction sites.

Distinct biogenesis and surveillance pathways are required to maintain the protective function of the multi-layered envelope of Gram-negative bacteria ^2^. Our CRISPRi analysis shows that DolP promotes efficient cell growth when the AmiA pathway of septal peptidoglycan splitting is impaired. This result is in line with the conclusions of a previous study implicating DolP in the regulation of NlpD-mediated activation of AmiC ^33^. Cells lacking both NlpD and AmiC have reduced OM integrity ^33^, which may contribute to explain the vancomycin sensitivity of Δ*dolP* cells. It remains poorly understood whether envelope stress caused by the accumulation of unfolded OMPs influences peptidoglycan remodelling. Intriguingly, our observation that mid-cell localization of DolP is reduced in *surA* and *bamB* deletion strains points to a possible role of DolP in linking envelope stress to septal peptidoglycan hydrolysis. Reduced levels of DolP at mid-cell sites, and thus impaired AmiC activation ^33^, could play an important role in coping with envelope stress, for instance by regulating the window of time available to restore normal OMP biogenesis prior to completing the formation of the new poles in the cell offspring.

Taken together, our results reveal an unprecedented link between activation of the envelope stress response and a late step of cell division. The fitness role of DolP during envelope stress emerges as an exploitable target in the development of new antibacterial therapies.

## Materials and Methods

### Bacterial strains and growth conditions

All *E. coli* strains used in this study are listed in Table S3. Strains newly generated for this study derive from BW25113 [Δ(*araD-araB*)*567* Δ(*rhaD-rhaB*)*568* Δ*lacZ4787* (::rrnB-3) *hsdR514 rph-1*] ^74^ or MG1655 (F^−^ λ^−^ *ilvG*^−^ *rfb-50 rph-1*) ^75^. Deletions of *dolP*, *rseA, surA, bamB, degP, or skp* in a BW25113 strain were achieved by P1 transduction of the Δ*dolP::kan^R^*, Δ*rseA::kan^R^*, Δ*surA::kan^R^*, Δ*bamB::kan^R^*, Δ*degP::kan^R^* or Δ*skp::kan^R^* alleles, respectively, obtained from the corresponding Keio collection strains ^37^. BW25113 derivative strains harbouring chromosomal fusions of constructs encoding superfolder GFP downstream of *dolP* or mCherry downstream of *nlpD*, *zipA* and *bamD* were obtained by λ-red recombination as previously described ^76^. Briefly, a kanamycin-resistance cassette was amplified from plasmid pKD4 using oligonucleotides carrying extensions of approximately 50 nucleotides homologous to regions immediately upstream or downstream the stop codon of the interested genes. *Dpn*I-digested and purified PCR products were electroporated into the BW25113 or derivative strains. Recombinant clones were selected at 37°C on LB agar plates containing kanamycin. When necessary, the *kan^R^* cassette inserted into a mutated locus (gene deletion or fusion) was removed upon transformation with the heat-curable plasmid pCP20 ^76^. The MG1655 derivative strain LC-E75, harbouring a dCas9-encoding construct under the control of the P_*tet*_ promoter, has been described ^38^. Cells were cultured in home-made lysogeny broth (LB) medium (1% (w/v) tryptone, 0.5% (w/v) yeast extract, 5 mg/ml (NaCl), commercially available Miller LB Broth (Sigma) or M9 minimal medium containing M9 salts (33.7 mM Na_2_HPO_4_, 22 mM KH_2_PO_4_, 8.55 mM NaCl, 9.35 mM NH_4_Cl) and supplemented with 0.2% w/v glycerol and all the amino acids, or all amino acids except methionine and cysteine in the case of ^35^S pulse-chase labeling. Antibiotics were used at the following concentrations: ampicillin 100 μg/ml, kanamycin 50 μg/ml, vancomycin 60 μg/ml. For spot tests, cells were cultured to mid-log phase, washed with M9 salts and serially diluted in ice-cold M9 salts prior to spotting on agar plates.

### Plasmid construction

All plasmids used in data figures are listed in Table S4. Plasmids for the ectopic expression of BAM subunits, DolP, or OmpA are derived from a pTrc99a vector. The plasmid pBAM^His^ (pJH114), which harbours a P_*trc*_ promoter followed by the *bamABCDE* open reading frames and an octahistidine tag fused downstream of *bamE*, was described ^55^. The region of pBAM^His^ comprising the segment that spans from the *bamA* start codon to the *bamE* stop codon was deleted by site-directed mutagenesis, generating pCtrl. Plasmids pBamA^His^ was generated by restriction-free cloning, inserting the *bamA* ORF without its stop codon downstream of the P_*trc*_ promoter and upstream of the octahistidine encoding region in pCtrl. pBamA^His^ was subjected to site directed mutagenesis to generate pBamA, encoding wild-type, non-tagged BamA. The *dolP* ORF amplified from the BW25113 genomic DNA was used to replace the *bamABCDE* ORFs in pJH114 by restriction-free cloning, generating pDolP^His^. The plasmid pBamA-DolP^His^ was generated by restriction-free cloning of the *bamA* ORF between the P_*trc*_ promoter and *dolP* in pDolP^His^. Site-directed mutagenesis was conducted on pBAM^His^ (pJH114) to obtain pBamACDE^His^, pBamABDE^His^, pBamBCDE^His^, pBamCDE^His^. A sequence encoding the tobacco etch virus protease cleavage site (TEV site) followed by a tandem Protein-A tag was amplified from pYM10 ^77^ and fused by restriction-free cloning with the last codon of the *bamE* gene in pBAM^His^ to generate pBAM^ProtA^. A stop codon was introduced downstream of the *bamE* last codon to generate pBAM. Plasmids encoding the ΔP1 and ΔP2 BamA variant were obtained by site-directed mutagenesis deleting the portion of *bamA* ORFs corresponding to residues E22-K89 or P92-G172, respectively. The TEV site and the tandem Protein-A construct amplified from pYM10 were inserted by restriction-free cloning downstream of the *dolP* last codon in pDolP^His^, generating pDolP^ProtA^. pDolP was derived from pDolP^ProtA^ using site-directed mutagenesis to introduce a stop codon immediately downstream of the *dolP* ORF. The *ompA* ORF was amplified from the BW25113 genomic DNA and inserted by restriction-free cloning between P_*trc*_ and the His-tag encoding construct of pCtrl to generate pOmpA^His^. The sgRNAs plasmids are derived from psgRNAcos ^38^. To generate sgRNA-encoding plasmids the DNA sequences AGCTGCACCTGCTGCGAATA (*bamD* sgRNA, plasmid pCAT187), GTAAACCACTCGCTCCAGAG (*bamE* sgRNA, plasmid pCAT189), CTCATCCGCGTGGGCGGAAA (*envC* sgRNA, plasmid pCAT191), and CTGAGCCGCCGACCGATTTA (*ftsX* sgRNA, pCAT193) were inserted into a *Bsa*I site of the psgRNAcos.

### CRISPRi screen and data analysis

Strain LC-E75 (*dolP*^+^) and its Δ*dolP* derivative were transformed with the EcoWG1 library which contains 5 guides per gene as previously described ^39^. After culturing pooled transformant cells in LB at 37°C to early exponential phase (optical density at 600 nm [OD_600_] = 0.2), a sample was withdrawn for plasmid isolation (t_start_). Subsequently, cultures were supplemented with 1 μM anhydrotetracycline (aTc) to induce dCas9 expression and further incubated at 37°C. When cultures reached an OD_600_ of 2 they were diluted 1:100 into LB supplemented with 1 μM aTc and incubated at the same temperature until an OD_600_ of 2. This step was repeated one more time prior to withdrawing a sample for isolation of plasmid DNA (t_end_). Sequencing indexes were used to assign reads to each sample. Illumina sequencing samples were prepared and analyzed as previously described ^38^. Briefly, a two-step PCR was performed with Phusion polymerase (Thermo Scientific) using indexed primers. The first PCR adds the first index and the second PCR adds the second index and flow-cell attachment sequences. Pooled PCR products were gel-purified. Sequencing was performed on a NextSeq550 machine (Illumina). The total number of reads obtained for each sample was used to normalize raw reads by sample size. Replicates were pooled to increase depth before another normalization by sample size. Guides with less than 100 normalized read counts in initial time points were discarded. For each screen, sgRNA fitness was calculated as the log2-transformed ratio of normalized reads counts between the final and the initial time point:

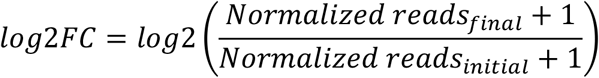

For each sample, log2FC values were centered by subtracting the median log2FC of non-targeting control guides. We then calculated for each sgRNA the difference of log2FC value between the Δ*dolP* screen and the *dolP*^+^ screen. Guides were ranked from the lowest negative values (negative fitness effect in Δ*dolP* compared to *dolP*^+^) to the highest positive values (positive fitness effect in Δ*dolP* compared to *dolP*^+^) and the significance of the interaction between *dolP* and each gene was evaluated by performing a minimum hypergeometric (mHG) test on the ranked list for each gene using the mHG R package (v. 1.1) ^78^. False-discovery rate (FDR) was used to correct p-values for multiple testing. For each gene, the median difference of log2FC between Δ*dolP* and *dolP*^+^ screens was used as a measure of the genetic interaction.

### Cell fractionation

To prepare whole-cell lysates, cells were cultured to early exponential phase (OD_600_ = 0.2-0.3) in LB medium at 37°C and collected. Where indicated, IPTG was added 60 min prior to cell collection. Cells were pelleted by centrifugation, washed once with M9 salt and lysed with Laemmli Sample Buffer (BioRad) (69 mM Tris-HCl, pH 6.8, 11.1% [v/v] glycerol, 1.1% [w/v] lithium dodecyl sulfate [LDS], 0.005% [w/v] bromophenol blue, supplemented with 357 mM β-mercaptoethanol and 2 mM phenylmethylsulfonyl fluoride [PMSF]). The total protein content was denatured at 98°C for 5 min prior SDS-PAGE analysis.

To perform lysozyme/EDTA lysis, cells were cultured to early exponential phase, collected by centrifugation, resuspended in 33 mM Tris-HCl pH 8 to an OD_600_ of 1. The cell suspension was then supplemented with 0.1 mg/ml lysozyme (Sigma), 2 mM EDTA and incubated on ice for 20 minutes to induce lysis. After addition of 10 mM MgSO_4_, the membrane fraction was collected by centrifugation at 16000x *g*. The supernatant was further centrifuged at 100000x *g* to remove any residual membrane fraction, which was discarded. The obtained soluble fraction was subjected to protein precipitation by adding 10% (w/v) trichloroacetic acid (TCA). TCA precipitates were solubilized in Laemmli Sample Buffer (BioRad) prior to SDS-PAGE analysis.

The crude envelope fractions directly analyzes by SDS-PAGE or used for native affinity purification of affinity tagged BAM complex or DolP were prepared from cells that were cultured in LB until early exponential phase and, where indicated, supplemented with 400 μM IPTG for 1 hour to induce ectopic expression. Cells were collected by centrifugation at 6000x *g* at 4°C, resuspended in 20 mM Tris-HCl pH 8, and mechanically disrupted using a Cell Disruptor (Constant Systems LTD) set to 0.82 kPa. The obtained cell lysate fractions were clarified by centrifugation at 6000x *g* and 4°C. The supernatant was then subjected to ultracentrifugation at 100000x *g* at 4°C to collect the envelope fraction.

### Isolation of native protein complexes by IgG- or nickel-affinity chromatography

The envelope fraction was resuspended at a concentration of approximately 10 mg/ml in solubilization buffer (20 mM Tris-HCl pH 7.4, 100 mM NaCl, 0.1 mM EDTA, 2 mM PMSF) supplemented with EDTA-free protease inhibitor cocktail (Roche), and 1.1% (w/v) of a mild detergent component corresponding to digitonin (Merck) and n-dodecyl β-D-maltoside (DDM, Merck) as indicated in Figure Legends. To facilitate extraction of membrane proteins, samples were subjected to mild agitation for 1 hour at 4°C. Insoluble material was removed by centrifugation at 16000x *g* at 4°C. To perform IgG affinity purification, membrane-extracted proteins were incubated for 1.5 hours at 4°C with purified human IgG (Sigma) that had been previously coupled to CNBr-activated Sepharose beads (GE Healthcare). After extensive washes of the resin with solubilization buffer containing 0.3% (w/v) digitonin and 0.03% (w/v) DDM, bound proteins were eluted by incubation with AcTEV protease (ThermoFisher) overnight at 4°C under mild agitation. To perform nickel (Ni)-affinity purification, membrane-extracted proteins were supplemented with 20 mM imidazole and incubated with Protino Ni-NTA agarose beads (Macherey-Nagel) for 1 hour at 4°C. After extensive washes of the resin with solubilization buffer supplemented with 50 mM imidazole, 0.3% (w/v) digitonin, 0.03% (w/v) DDM, and the EDTA-free protease inhibitor cocktail (Roche), bound proteins were eluted using the same buffer supplemented with 500 mM imidazole.

### *In vitro* reconstitution of the BAM-DolP interaction and BN-PAGE analysis

Envelope fractions were obtained from cells carrying pBAM^His^ or pDolP^His^ and cultured until early exponential phase in LB medium at 37°C and subsequently supplemented with 400 μM IPTG for 1.5 hours to induce the expression of the BAM complex genes or *dolP*. The envelope fractions were solubilized and purified by Ni-affinity and size exclusion chromatography, adapting a previously published protocol. Briefly, after membrane solubilization with 50 mM Tris-HCl pH 8.0, 150 mM NaCl, and 1% (w/v) DDM, and removal of insoluble material by ultracentrifugation at 100000x *g*, 4°C, soluble proteins were loaded onto a Ni-column (HisTrap FF Crude, GE Healthcare) pre-equilibrated with 50 mM Tris-HCl pH 8.0, 150 mM NaCl, and 0.03% (w/v) DDM (equilibration buffer), using an ÄKTA Purifier 10 (GE Healthcare) at 4°C. The column containing bound proteins was washed with equilibration buffer supplemented with 50 mM imidazole. Proteins were eluted in equilibration buffer, applying a gradient of imidazole from 50 mM to 500 mM and further separated by gel filtration using an HiLoad 16/600 Superdex 200 (GE Healthcare) in equilibration buffer. Eluted proteins were concentrated using an ultrafiltration membrane with a 10 kDa molecular weight cutoff (Vivaspin 6, Sartorius). To reconstitute the BAM-DolP complex *in vitro*, equimolar concentrations of purified BAM and DolP were used. Purified proteins were mixed in equilibration buffer for 1 hour at 4°C or for 30 min at 25°C as indicated in Figure Legends. The reaction was further diluted 1:4 times in ice-cold blue native buffer (20 mM Tris-HCl pH 7.4, 50 mM NaCl, 0.1 mM EDTA, 1% [w/v] digitonin, 10% w/v glycerol) and ice cold blue native loading buffer (5% coomassie brilliant blue G-250, 100 mM Bis-Tris-HCl, pH 7.0, 500 mM 6-aminocaproic acid) prior to loading onto home-made 5-13% blue native polyacrylamide gradient gels. Resolved protein complexes were blotted onto a PVDF membrane and immunolabeled. Where non-relevant gel lanes were removed, a white space was used to separate contiguous parts of the same gel.

### Pulse-chase *in vivo* protein labeling

Pulse-chase labeling of cellular proteins with radioactive ^35^S methionine and cysteine was conducted as previously described ^79^. Briefly, overnight cultures were washed and freshly diluted into M9 medium to OD_600_ = 0.03 and incubated at 37 °C. When the cultures reached OD_600_ = 0.3, ectopic expression of EspP was induced by adding 200 μM IPTG for 30 min. ^35^S labelling was performed by supplementing cultures with a mixture of [^35^S]-methionine and [^35^S]-cysteine followed after 30 seconds by the addition of an excess of cold methionine and cysteine. Samples were withdrawn at different chase times, immediately mixed with an equal volume of ice and subjected to TCA precipitation, prior to immunoprecipitation.

### Antibodies

Proteins separated by SDS-PAGE or blue native-PAGE and blotted onto PVDF membranes were immunodecorated using epitope-specific rabbit polyclonal antisera with the following exceptions. The F_1_β subunit of the ATP F_1_F_0_ synthase was detected using a rabbit polyclonal antiserum raised against an epitope of the homologous protein of *Saccharomyces cerevisiae* (Atp2). RpoB was detected using a mouse monoclonal antibody (NeoClone Biotechnology). The secondary immunodecoration was conducted using anti-rabbit or anti-mouse peroxidase-conjugated antibodies produced in goat (Sigma). EspP immunoprecipitation was conducted using a rabbit polyclonal antiserum specific for a C-terminal epitope of EspP.

### Epifluorescence microscopy and analysis

Overnight cultures of *E. coli* BW25113 and its derivative strains were diluted into fresh M9 medium containing 0.2% glycerol or LB medium and grown at 30°C to OD_600_ = 0.2-0.3. When indicated, cultures were supplemented with 400 μM of IPTG to induce ectopic expression of plasmid-borne genes for 1 hour prior to collecting samples for microscopy analysis. Culture volumes of 0.6 μl were deposited directly onto slides coated with 1% (w/v) agarose in a phosphate-buffered saline solution and visualized by epifluorescence microscopy. Cells were imaged at 30°C using an Eclipse TI-E/B Nikon wide field epifluorescence inverted microscope with a phase contrast objective (Plan APO LBDA 100X oil NA1.4 Phase) and a Semrock filter mCherry (Ex: 562BP24; DM: 593; Em: 641BP75) or FITC (Ex: 482BP35; DM: 506; Em: 536BP40). Images were acquired using a CDD OrcaR2 (Hamamatsu) camera with illumination at 100% from a HG Intensilight source and with an exposure time of 1-3 seconds, or using a Neo 5.5 sCMOS (Andor) camera with illumination at 60% from a LED SPECTRA X source (Lumencor) with an exposure time of 2 seconds. Nis-Elements AR software (Nikon) was used for image capture. Image analysis was conducted using the Fiji and ImageJ software. The fraction of cells with DolP^GPF^ signals at mid-cell sites was estimated using the Fiji Cell Counter plugin. Collective profiles of fluorescence distribution versus the relative position along the cell axis were generated using the Coli-Inspector macro run in ImageJ within the plugin ObjectJ ^80^, selecting only cells with a constriction (80% of cell diameter) as qualified objects.

## Acknowledgments

We thank Harris Bernstein (NIDDK/NIH, Bethesda, MD) for providing key reagents and for comments on the manuscript. We thank Tanneke Den Blaauwen (University of Amsterdam) and Nathalie Dautin (IBPC/CNRS, Paris) for discussion. We thank the Light Imaging Toulouse CBI (LITC) platform for assistance with and maintenance of the microscopy instrumentation. Financial support was provided by: the Fondation pour la Recherche Médicale to D.R.; the Chinese Scholarship Council to Y.Y.; the Ecole Normale Supérieure to F.R.; the European Research Council (Europe Union’s Horizon 2020 research and innovation program, grant agreement No 677823), the French governmental Investissement d’Avenir program and the Laboratoire d’Excellence “Integrative Biology of Emerging Infectious Diseases” (ANR-10-LABX-62-IBEID) to D.B.; the CNRS ATIP-Avenir program to R.I.

## Author contribution

R.I. conceived the project and wrote the manuscript; D.B. conceived and developed the CRISPRi approach; R.I., D.B., D.R. L.C. and C.A. supervised the experiments; D.R., Y.Y., L.O., F.R., L.C., V.M., C.M., G.M., A.C.-S, C.T., J.R. and C.A. performed the experiments and analyzed the data together with R.I. and D.B.; A.C.-S, D.R., Y.Y., L.O. and F.R. prepared the figures; all authors discussed the project and the experimental results, and commented on the manuscript.

---

**Fig. S1.**
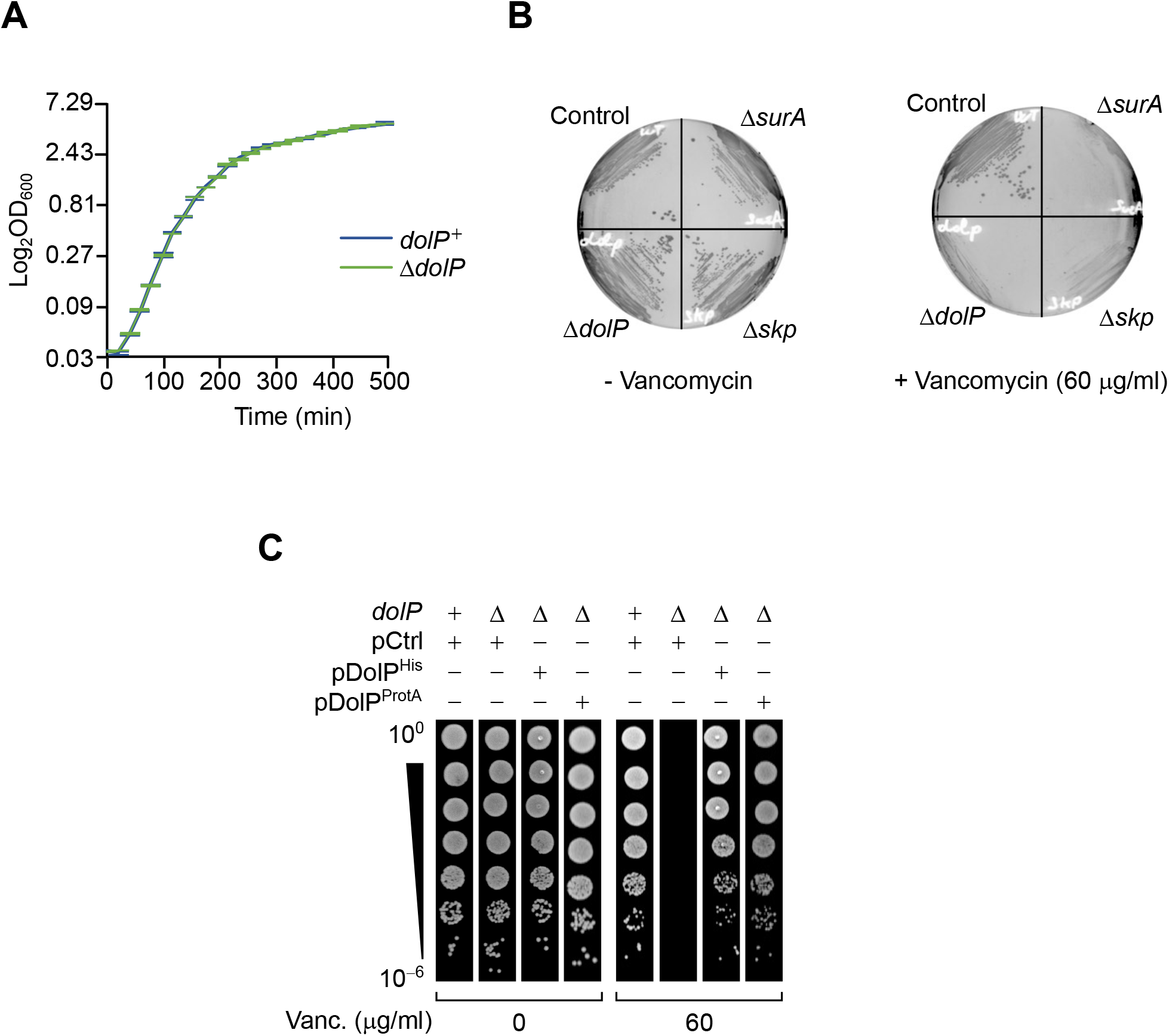
The deletion of *dolP* severely impairs growth in the presence of vancomycin. (**A**) BW25113 and Δ*dolP* cells were cultured in LB. The cell densities of both cultures were monitored by measuring the OD_600_ at regular time intervals. The graph reports mean values of independent cultures ± standard deviation (SD, *n* = 3). (**B**) BW25113 (control) and the indicated deletion strains were cultured on LB agar lacking or containing 60 μg/ml vancomycin. (**C**) BW25113 and Δ*dolP* cells carrying the indicated plasmids were serially diluted and spotted on LB lacking or containing 60 μg/ml vancomycin. Ectopic expression was driven by the leaky transcriptional activity of P_*trc*_ in the absence of IPTG.

**Fig. S2.**
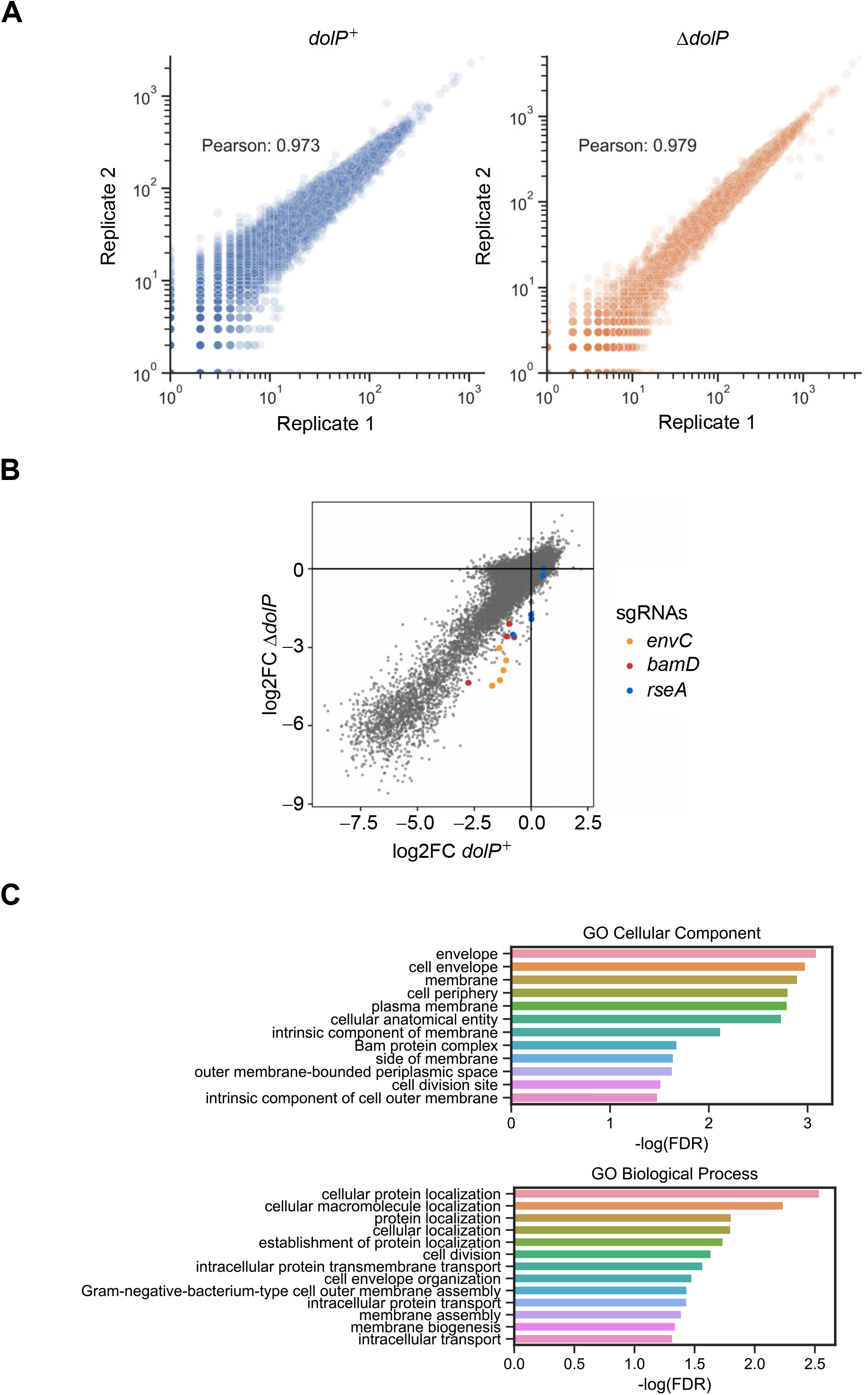
Reproducibility of the CRISPRi screens and ontology analysis of gene hits. (**A**) The raw read counts of experimental replicates are very well correlated (Pearson’s r > 0.97) in both *dolP*^+^ (left) and Δ*dolP* (right) screens. (**B**) Log2FC values of each sgRNA are compared between the WT and Δ*dolP* screens. Only guides targeting *envC*, *bamD*, and *rseA* are highlighted in orange, red and cyan, respectively. (**C**) Gene Ontology analysis of the genes with a significant synthetic interaction with *dolP*. Using FDR<0.05 as a threshold, 27 genes were selected for Gene Ontology (GO) analysis for cellular component (upper panel) or biological process (bottom panel) ^81^. Both analyses show an overrepresentation of genes associated with cell envelope and membrane function.

**Fig. S3.**
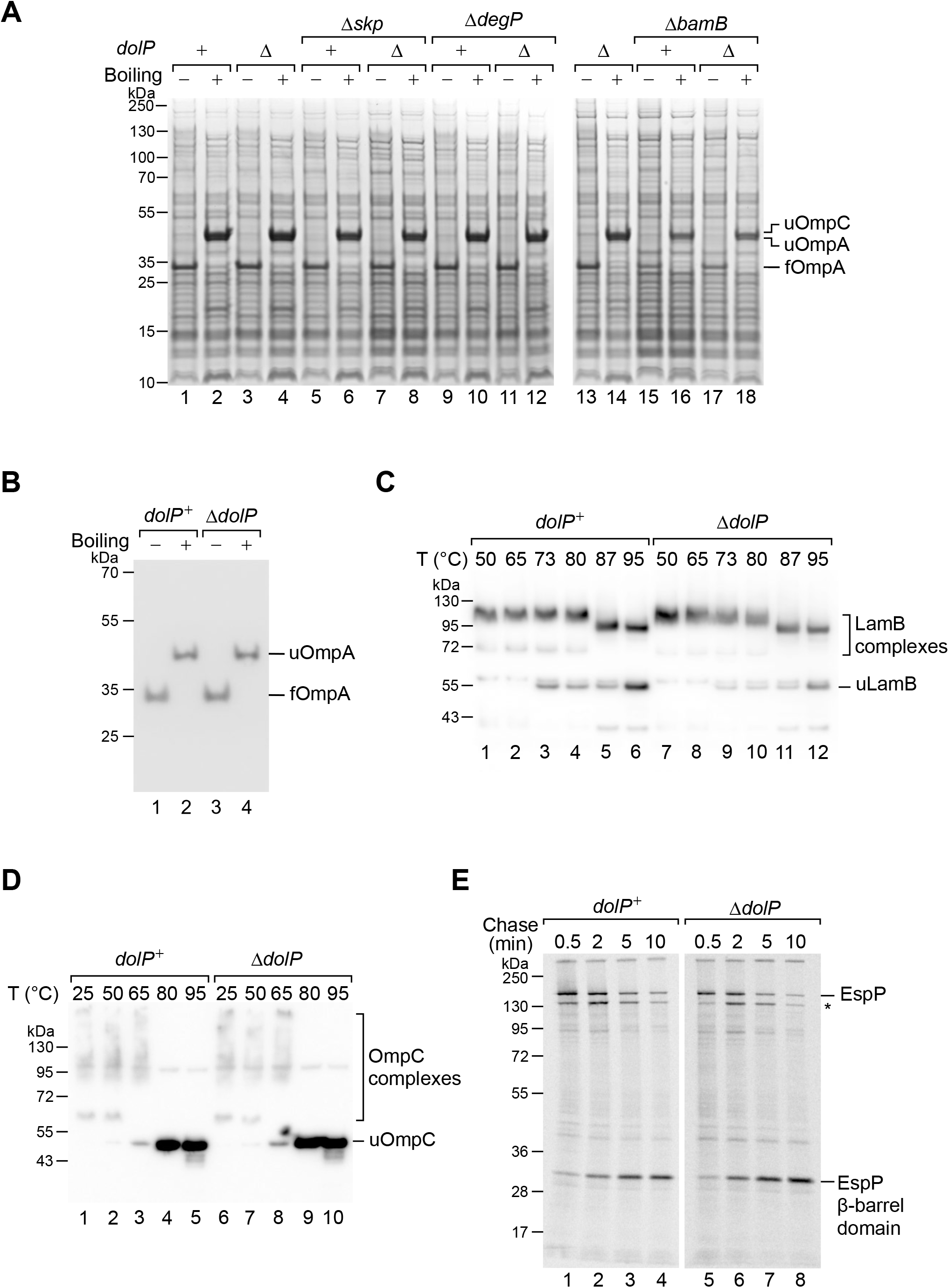
DolP is not crucial for OMP biogenesis. (**A**) The protein contents of the indicated envelope fractions were analysed by SDS-PAGE and coomassie staining. Prior to loading, samples were heated at 50°C (boiling −) or 99°C (boiling +) for 10 min. (**B**) The protein contents of the envelope fraction of BW25113 (*dolP*^+^) or Δ*dolP* cells were mixed with SDS-PAGE loading buffer, heated at 50°C (boiling −) or 99°C (boiling +) for 10 min, and analyzed by SDS-PAGE and immunoblotting. Unfolded (u) and folded (f) OmpA forms are indicated. (**C** and **D**) The protein contents of the envelope fraction of BW25113 (*dolP*^+^) and Δ*dolP* cells was mixed with SDS-PAGE loading buffer, heated at the indicated temperatures, separated by SDS-PAGE and immunoblotted using anti-LamB (C) or anti-OmpC (D) antisera. LamB and OmpC form complexes (trimers), which are resistant to SDS and require significant heating to be denatured. At progressively higher temperatures, LamB (C) and OmpC (D) complexes are denatured and the two proteins migrate as monomers (u, unfolded). (**E**) The biogenesis of EspP, an autotransporter OMP of enterohemorrhagic *E. coli* heterologously expressed in BW25113 cells, was monitored in time. The plasmid pRI22 ^79^ harbouring the EspP encoding gene under the control of an IPTG-inducible promoter was transformed into wild-type BW25113 (*dolP*^+^) and Δ*dolP* cells. EspP is a serine protease autotransporter of *Enterobacteriaceae*, and consists of an N-terminal passenger domain secreted on the cell surface and a C-terminal domain which is ultimately folded into an integral OM β-barrel structure ^82^. EspP biogenesis requires both periplasmic chaperones, such as SurA, and the BAM complex ^79^. Once EspP assembly and secretion are completed, the passenger domain is cleaved by an autocatalytic reaction that is catalyzed by residues of the folded β-barrel domain facing the interior of the barrel. Thus, the proteolytic reaction can be exploited to infer about completion of β-barrel folding and assembly ^82^. To monitor the kinetics of EspP autoproteolysis, cells carrying pRI22 were cultured in M9 minimal medium until early exponential phase, supplemented with 200 μM IPTG and subjected to ^35^S pulse-chase labelling. The total protein contents of samples collected after the indicated chase times were subjected to immunoprecipitation using an antiserum specific for an EspP C-terminal peptide. The asterisk indicates a degradation product of EspP, likely generated by the OM protease OmpT, which cleaves the N-terminal region of the passenger domain as it is secreted across the OM^83^. A band corresponding to the EspP β-barrel domain is generated with a similar time dependence in both *dolP*^+^ and Δ*dolP* cells.

**Fig. S4.**
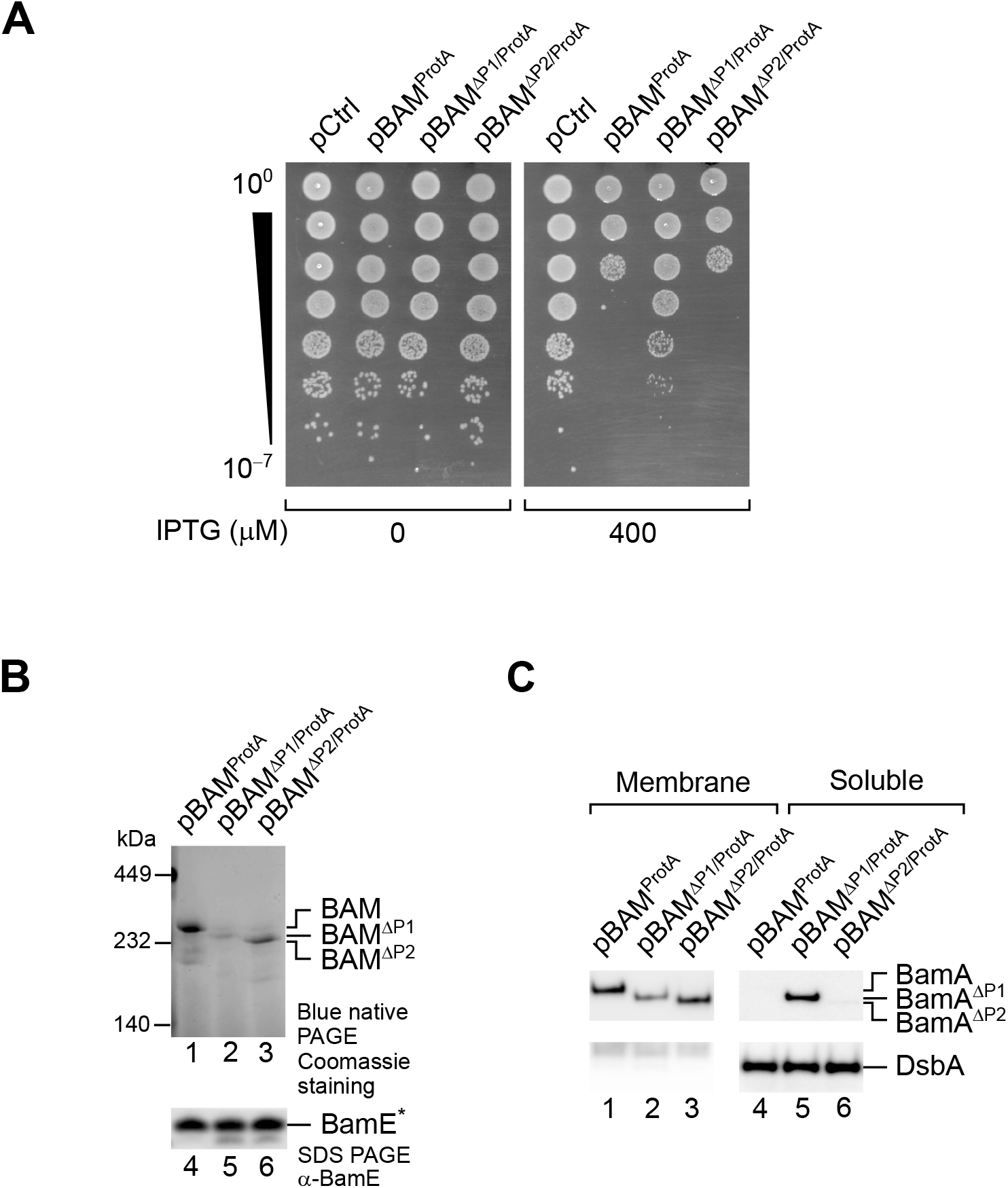
The detrimental effect of BAM overproduction is caused by the overaccumulation of BAM in the OM. (**A**) BW25113 cells were transformed with an empty vector or vectors for the ectopic expression of the BAM complex containing protein A-tagged BamE (pBAM^ProtA^), or similar variants of the BAM complex including BamA^ΔP1^ (pBAM^ΔP1/ProtA^, lacking the N-terminal POTRA1 motif) or BamA^ΔP2^ (pBAM^ΔP2/ProtA^, lacking the POTRA2 motif) in place of wild-type BamA. In the presence of IPTG, overproduction of BAM^ΔP1/ProtA^ impaired growth to a lower extent compared to overproduction of BAM^ProtA^ or BAM^ΔP2/ProtA^. (**B**) The crude envelope fractions of cells overproducing BAM^ProtA^, BAM^ΔP1/ProtA^, or BAM^ΔP2/ProtA^ were solubilized with 1% (w/v) digitonin and 0.1% (w/v) DDM, and subjected to native IgG-affinity chromatography. Elutions were analyzed by blue native-PAGE and coomassie staining (lanes 1-3) or SDS-PAGE and immunoblotting (lanes 4-6). Blue native-PAGE of the elution fraction containing wild-type BamA resolved a coomassie stainable complex migrating with an apparent mass of 250 kDa, which corresponds to the BAM complex (lane 1). A roughly similar amount of BAM complex was detected in the elution obtained with BamA^ΔP2/ProtA^ samples, although in this case the BAM^ΔP2^ complex migrated slightly faster, accounting for the BamA^ΔP2^ mass difference (lane 3). In contrast, the amount of BAM^ΔP1/ProtA^ variant was considerably lower (lane 2). The asterisk indicates that BamE is obtained from TEV digestion of BamE^ProtA^. (**C**) Cells overproducing the indicated variants of the BAM complex were fractionated. The membrane and soluble fractions, obtained upon treatment of collected cells with lysozyme and EDTA were analyzed by SDS-PAGE and immunoblotting. BamA^ΔP1^ was depleted from the total membrane fraction (lanes 1-3), and accumulated in the soluble fraction containing the periplasmic protein DsbA (lanes 4-6).

**Fig. S5.**
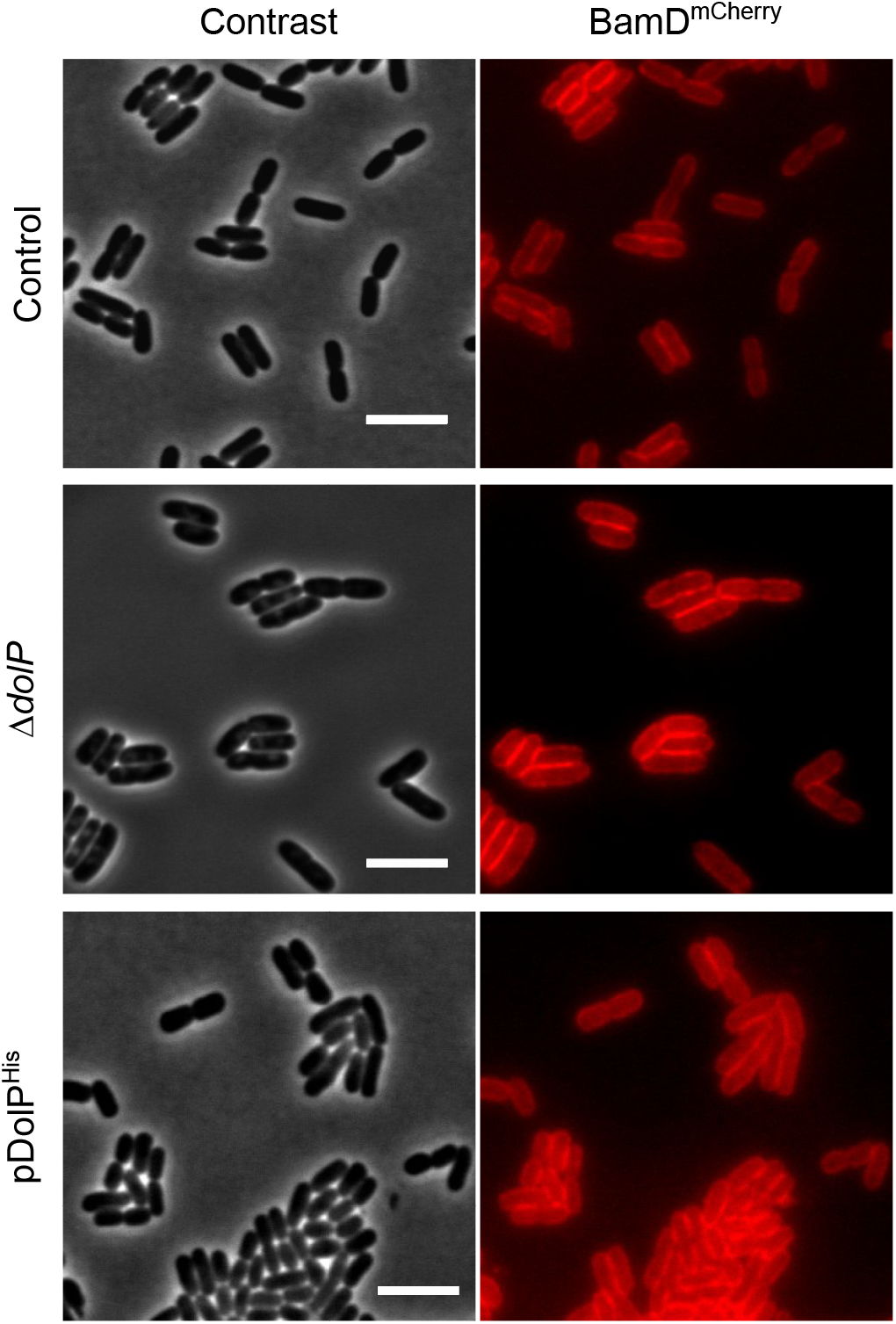
Effect of lack or overproduction of DolP on BAM localization. BW25113 derivative cells harbouring a *bamD-mCherry* chromosomal fusion were visualized on 1% (w/v) agarose pads by fluorescence and phase-contrast microscopy. Top: *dolP*^+^ cells (control); Center: Δ*dolP* cells: Bottom: *dolP*^+^ cells carrying pDolP^His^. Cells were cultured at 30°C in minimal M9 medium to OD_600_ = 0.5. IPTG (400 μM) was supplemented for 1 hour to induce ectopic expression of DolP^His^. Cells were visualized on 1% (w/v) agarose pads by fluorescence and phase-contrast microscopy. Bar = 5 μm.

**Fig. S6.**
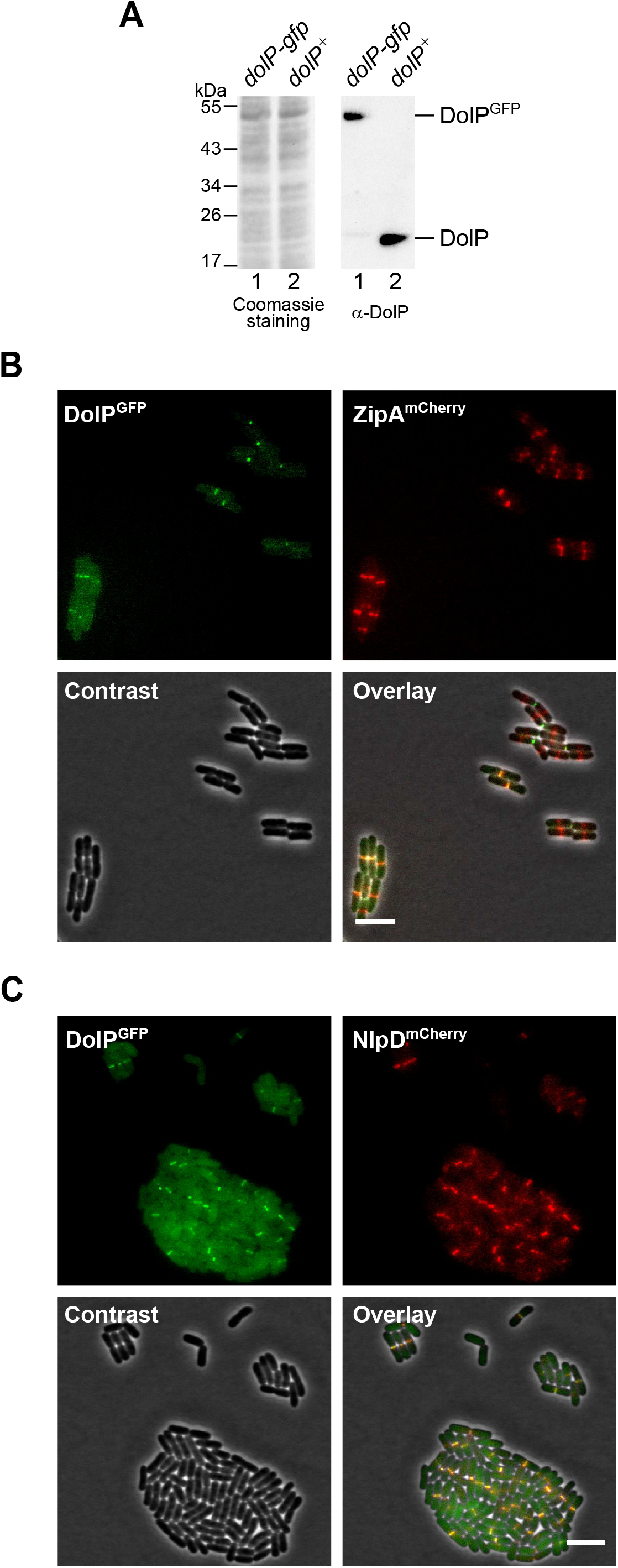
DolP accumulates at mid-cell sites during a late step of cell division. (**A**) The total protein content of cells harbouring the *dolP-gfp* chromosomal fusion (lanes 1) or wild-type *dolP* (lanes 2) were analyzed by SDS-PAGE followed by coomassie staining or immunoblotting using a DolP specific antiserum. (**B, C**) BW25113-derivative cells harbouring the chromosomal fusion *dolP-gfp* and either *zipA-mCherry* (B) or *nlpD-mCherry* (C) were cultured at 30°C in minimal M9 medium to OD_600_ = 0.2-0.3 and visualized on 1% (w/v) agarose pads by fluorescence and phase-contrast microscopy. Bar = 5 μm.

**Fig. S7.**
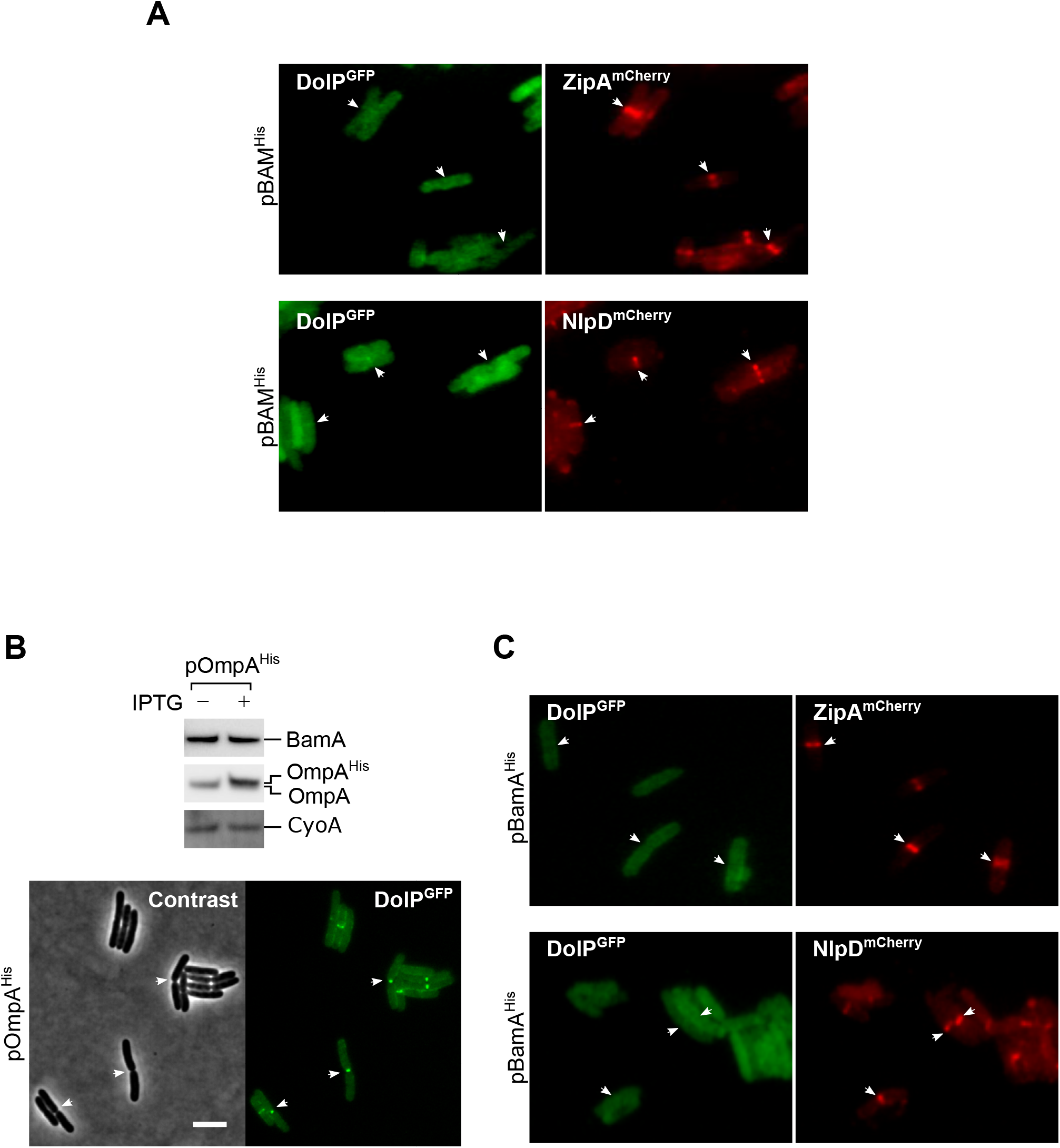
Overproduction of BAM does not influence septal recruitment of NlpD and ZipA. (**A**) BW25113-derivative cells harbouring the chromosomal fusion *dolP-gfp* and *zipA-mCherry* or *nlpD-mCherry* were transformed with pBAM^His^ (ectopic overproduction of all BAM subunits, with His-tagged BamE). Cells were then cultured at 30°C in minimal M9 medium supplemented with 400 μM IPTG for 1 hour until OD_600_ = 0.2-0.3, and visualized on 1% (w/v) agarose pads by fluorescence microscopy. The arrowheads indicate the localization of DolP^GFP^, ZipA^mCherry^ or NlpD^mCherry^ at division septa. (**B**) Top: BW25113-derivative *dolP-gfp* cells transformed with pOmpA (ectopic overproduction of His-tagged OmpA) were culture in LB medium and supplemented with 400 μM IPTG or no IPTG for 1 hour prior to collecting cells. The total protein content was analyzed by SDS-PAGE and immunoblotting using the indicated antisera. Bottom: BW25113-derivative *dolP-gfp* cells transformed with pOmpA^His^ were cultured in minimal M9 medium supplemented with 400 μM IPTG for 1 hour until OD_600_ = 0.2-0.3, and visualized (A). Bar = 5 μm. (**C**) BW25113-derivative cells harbouring the chromosomal fusion *dolP-gfp* and *zipA-mCherry* or *nlpD-mCherry* were transformed with pBamA^His^ (ectopic overproduction of a partially inactive His-tagged form of BamA). Cells were then cultured at 30°C in minimal M9 medium supplemented with 400 μM IPTG for 1 hour until OD_600_ = 0.2-0.3, and visualized as in (A).

**Fig. S8.**
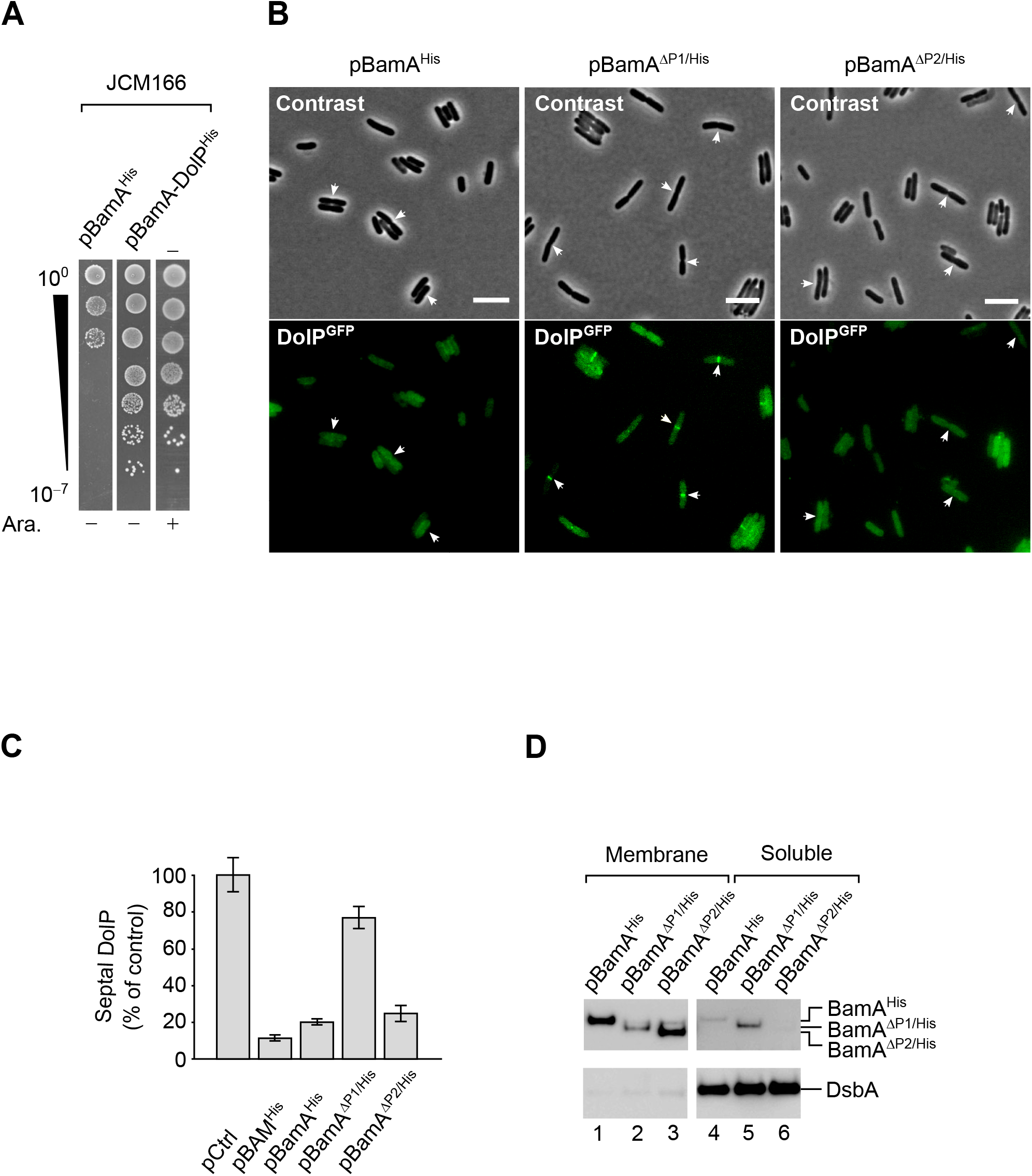
BamA overaccumulation in the OM impairs septal DolP localization. (**A**) The JCM166 BamA depletion strain or its transformants carrying plasmids encoding BamA^His^ or wild-type BamA and DolP^His^ were cultured in the presence of 0.02% (w/v) arabinose (which induces transcription of a chromosomal copy of wild-type *bamA* engineered downstream of an arabinose-inducible P_*bad*_ promoter). Serial dilutions were spotted on LB agar devoid of or supplemented with 0.02% (w/v) arabinose, as indicated. In the absence of arabinose, growth is supported by the ectopic expression of wild-type BamA and not of its variant encoding C-terminally His-tagged BamA. (**B**) Overnight cultures of BW25113 cells harboring the chromosomal *dolP-gfp* fusion and carrying the indicated plasmids were diluted in minimal M9 medium supplemented with 400 μM IPTG. The cells were grown at 30°C to OD_600_ = 0.2-0.3 and visualized on 1% (w/v) agarose pads by fluorescence microscopy. The arrows indicate envelope constriction sites between forming daughter cells. Bar = 5 μm. (**C**) Counting of BW25113 cells presenting septal DolP^GFP^ fluorescent signals. *dolP-gfp* cells carring pBAM^His^, pBamA^His^, pBamA^ΔP1/His^ or pBamA^ΔP2/His^ (analyzed by fluorescence microscopy and shown in Fig. 5A and in Fig. S8B) were counted. The percentage of cells presenting fluorescent DolP^GFP^ signals at the mid-cell in samples overproducing all five BAM subunits or only one of the indicated BamA variants was normalized to the same fraction obtained for cells carrying the control empty vector (pCtrl). Bar charts display a mean value ± SD (*n* = 3). More than 300 cells were counted in each experiment. (**D**) BW25113 cells carrying the indicated plasmids were cultured and supplemented with 400 μM IPTG to induce ectopic expression of BamA^His^ and its mutant forms. Collected cells were subjected to lysozyme/EDTA lysis to obtain the total membrane and soluble fractions. The protein contents of the indicated cell fractions were separated by SDS-PAGE and immunolabeled with the indicated antisera.

**Fig. S9.**
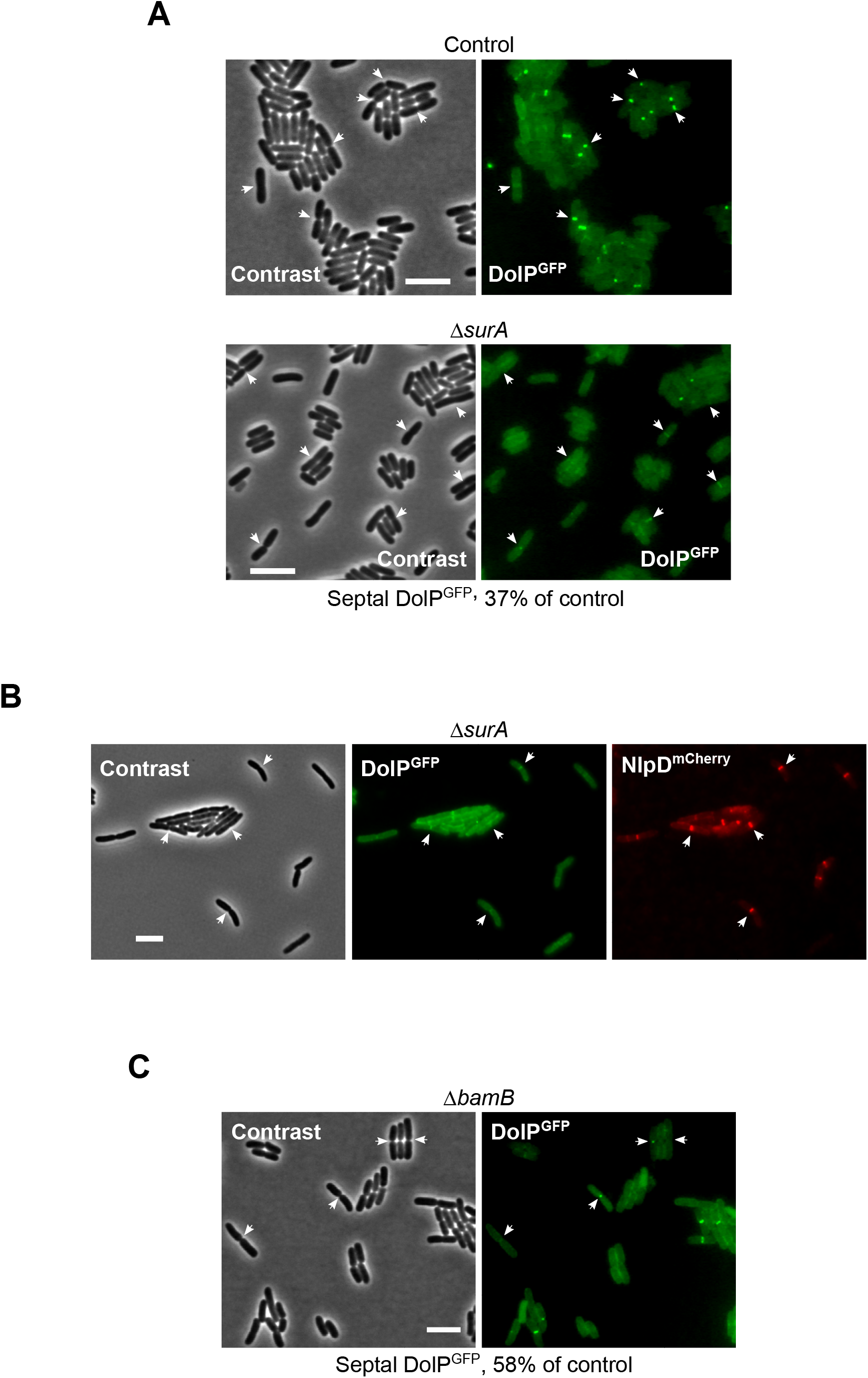
Envelope stress influences DolP localization. (**A**) Overnight cultures of BW25113 (Control) or Δ*surA* derivative cells carrying the *dolP-gfp* chromosomal fusion were freshly diluted in M9 medium and incubated at 30°C till OD_600_ = 0.03 and visualized on 1% (w/v) agarose pads by contrast and fluorescence microscopy. Bar = 5 μm. The arrows indicate envelope constriction sites between forming daughter cells. The percentage of cells presenting fluorescent DolP^GFP^ signals at division septa in Δ*surA* cells was normalized to the same fraction obtained for the control cells and reported below the micrograph. More than 1000 cells were counted in each sample. (**B**) Overnight cultures of BW25113 (control) or Δ*surA* derivative cells carrying the *dolP-gfp* and *nlpD*-*mCherry* chromosomal fusions were freshly diluted in LB medium and incubated at 30°C until OD_600_ = 0.3 prior to visualization by contrast and fluorescence microscopy. Bar = 5 μm. The arrows indicate envelope constriction sites between forming daughter cells (**C**) Overnight cultures of Δ*bamB* cells harboring the chromosomal *dolP-gfp* fusion were diluted in minimal M9 medium, cultured at 30°C to OD_600_ = 0.3 and visualized on 1% (w/v) agarose pads by contrast and fluorescence microscopy. The arrows indicate envelope constriction sites between forming daughter cells. Bar = 5 μm. The percentage of cells presenting fluorescent DolP^GFP^ signals at division septa in Δ*bamB* cells was normalized to the same fraction obtained for the control cells (BW25113 carrying the *dolP-gfp* chromosomal fusion) and reported below the micrograph. More than 300 cells were counted.

**Table S1. Log2FC values of sgRNAs in the WT or Δ *dolP* screen.**

See source data file (Table S1)

**Table S2. Genetic interaction scores.**

See source data file (Table S2)

**Table S3:** List of strains.

| Name | Genotype and relevant features | Source |
| --- | --- | --- |
| BW25113 | $\Delta(araD-araB)567 \Delta(rhaD-rhaB)568 \Delta lacZ4787(::rrnB-3) hsdR514 rph-1$ (wild-type reference) | Grenier *et al.*, 2014 <sup>74</sup> |
| $\Delta rseA$ | BW25113 *rseA::kan* | This study |
| $\Delta dolP$ | BW25113 *dolP::kan* | This study |
| $\Delta bamB$ | BW25113 *bamB::kan* | This study |
| $\Delta bamB \Delta dolP$ | BW25113 $\Delta bamB dolP::kan$ | This study |
| $\Delta dolP \Delta rseA$ | BW25113 $\Delta dolP rseA::kan$ | This study |
| $\Delta skp$ | BW25113 *skp::kan* | This study |
| $\Delta dolP \Delta skp$ | BW25113 $\Delta dolP skp::kan$ | This study |
| $\Delta degP$ | BW25113 *degP::kan* | This study |
| $\Delta dolP \Delta degP$ | BW25113 $\Delta dolP degP::kan$ | This study |
| *dolP-gfp* | BW25113 *dolP-gfp* | This study |
| $\Delta surA$ | BW25113 *surA::kan* | This study |
| $\Delta surA dolP-gfp$ | BW25113 *surA::kan dolP-gfp* | This study |
| $\Delta bamB dolP-gfp$ | BW25113 *bamB::kan dolP-gfp* | This study |
| $\Delta dolP bamD-mCherry$ | BW25113 $\Delta dolP bamD-mCherry$ | This study |
| *bamD-mCherry* | BW25113 *bamD-mCherry* | This study |
| *dolP-gfp nlpD-mCherry* | BW25113 *dolP-gfp nlpD-mCherry* | This study |
| $\Delta surA dolP-gfp nlpD-mCherry$ | BW25113 *dolP-gfp nlpD-mCherry surA::kan* | This study |
| *dolP-gfp zipA-mCherry* | BW25113 *dolP-gfp zipA-mCherry* | This study |
| JCM166 | MC4100 *ara<sup>r/-</sup> Δ(λatt-lom)::bla P<sub>BAD</sub>yaeT araC ΔyaeT* (BamA depletion strain) | Wu *et al.*, 2005 <sup>19</sup> |
| LC-E75 | $F^- \lambda^- ilvG^- rfb-50 rph-1 186attB::P_{tet^-}dcas9, \lambda attB::mCherry$ | Cui *et al.*, 2018 <sup>38</sup> |
| LC-E75 $\Delta dolP$ | LC-E75 *dolP::kan* | This study |

**Table S4:**
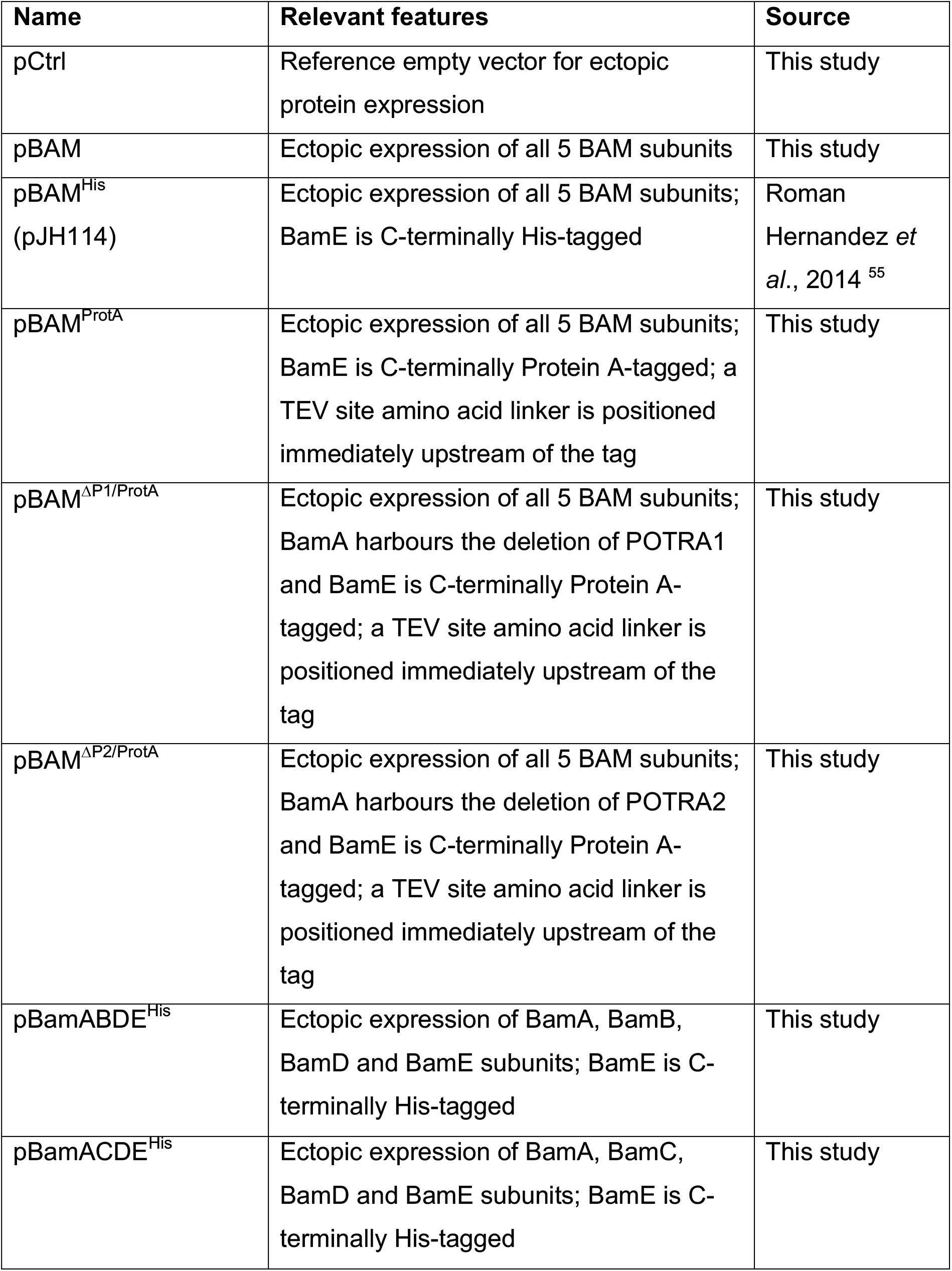
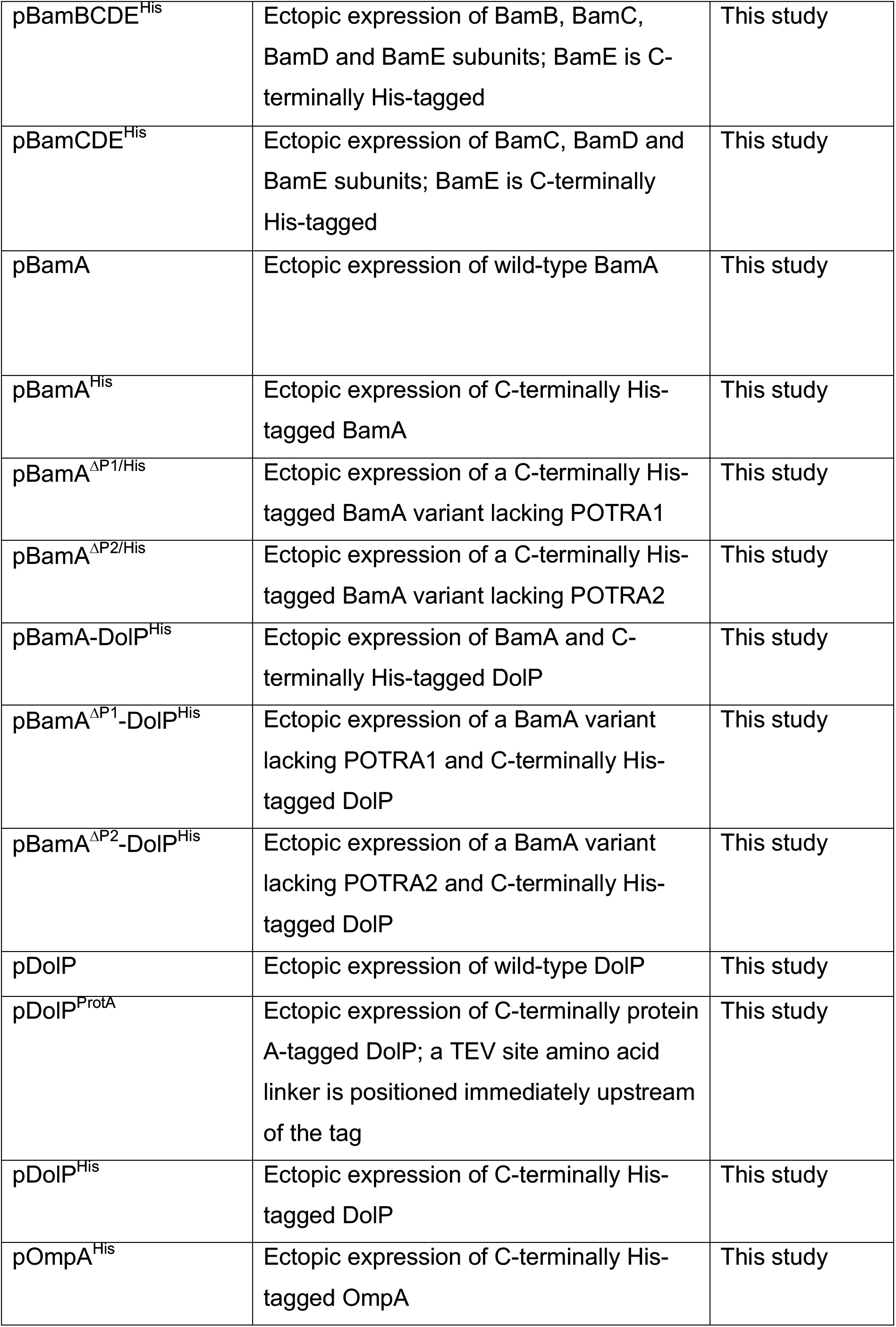
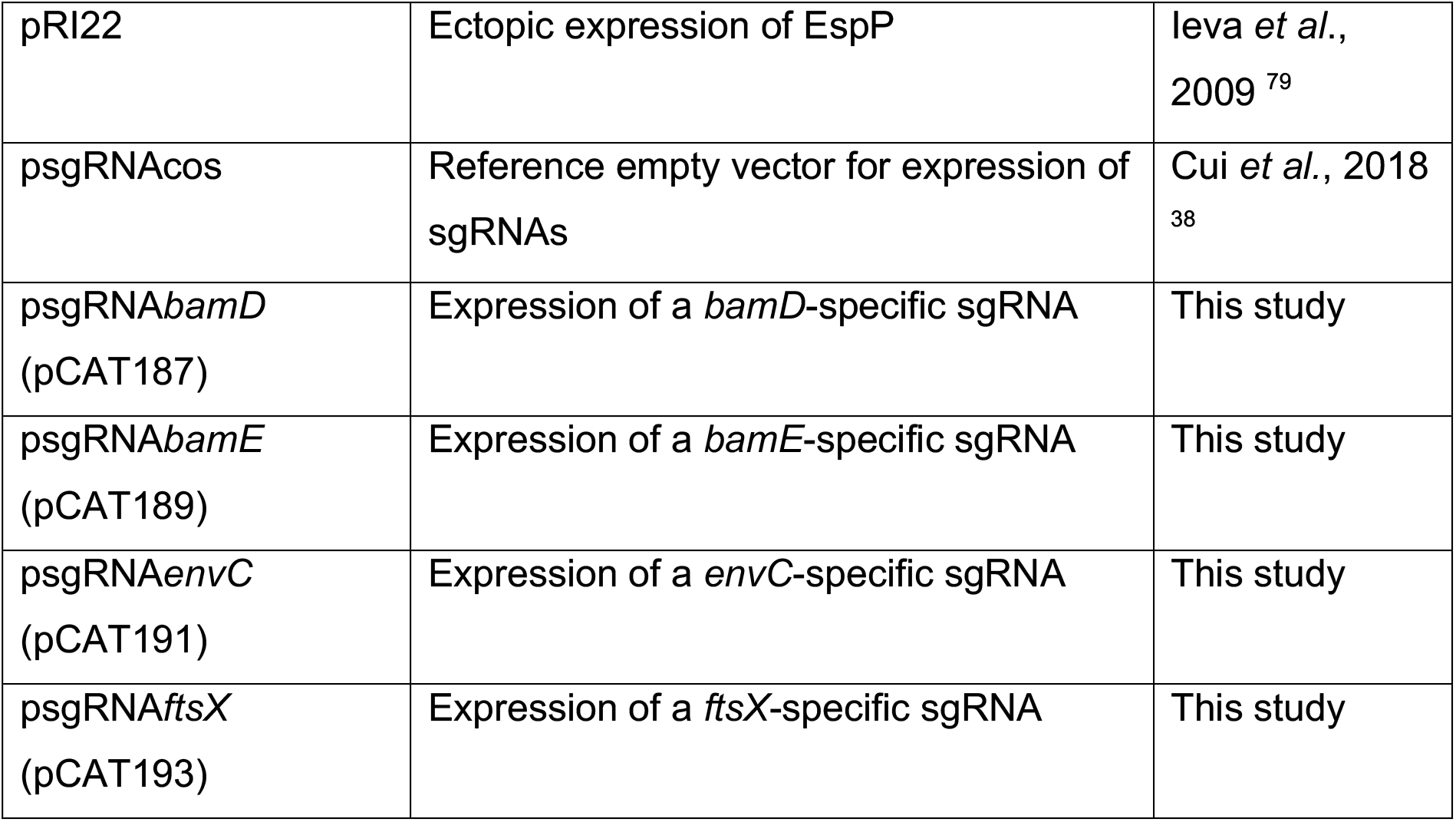
List of plasmids.

## References

1 den Blaauwen, T., Hamoen, L. W. & Levin, P. A. The divisome at 25: the road ahead. Current opinion in microbiology 36, 85–94, doi:10.1016/j.mib.2017.01.007 (2017).

2 Egan, A. J. F., Errington, J. & Vollmer, W. Regulation of peptidoglycan synthesis and remodelling. Nature reviews. Microbiology, doi:10.1038/s41579-020-0366-3 (2020).

3 Calmettes, C., Judd, A. & Moraes, T. F. Structural Aspects of Bacterial Outer Membrane Protein Assembly. Advances in experimental medicine and biology 883, 255–270, doi:10.1007/978-3-319-23603-2_14 (2015).

4 Nikaido, H. Molecular basis of bacterial outer membrane permeability revisited. Microbiology and molecular biology reviews : MMBR 67, 593–656, doi:10.1128/mmbr.67.4.593-656.2003 (2003).

5 Ranava, D., Caumont-Sarcos, A., Albenne, C. & Ieva, R. Bacterial machineries for the assembly of membrane-embedded beta-barrel proteins. FEMS microbiology letters 365, doi:10.1093/femsle/fny087 (2018).

6 Schiffrin, B., Brockwell, D. J. & Radford, S. E. Outer membrane protein folding from an energy landscape perspective. BMC biology 15, 123, doi:10.1186/s12915-017-0464-5 (2017).

7 Webb, C. T., Heinz, E. & Lithgow, T. Evolution of the beta-barrel assembly machinery. Trends in microbiology 20, 612–620, doi:10.1016/j.tim.2012.08.006 (2012).

8 Voulhoux, R., Bos, M. P., Geurtsen, J., Mols, M. & Tommassen, J. Role of a highly conserved bacterial protein in outer membrane protein assembly. Science (New York, N.Y.) 299, 262–265, doi:10.1126/science.1078973 (2003).

9 Noinaj, N., Kuszak, A. J., Gumbart, J. C., Lukacik, P., Chang, H., Easley, N. C., Lithgow, T. & Buchanan, S. K. Structural insight into the biogenesis of beta-barrel membrane proteins. Nature 501, 385–390, doi:10.1038/nature12521 (2013).

10 Iadanza, M. G., Higgins, A. J., Schiffrin, B., Calabrese, A. N., Brockwell, D. J., Ashcroft, A. E., Radford, S. E. & Ranson, N. A. Lateral opening in the intact beta-barrel assembly machinery captured by cryo-EM. Nature communications 7, 12865, doi:10.1038/ncomms12865 (2016).

11 Gu, Y., Li, H., Dong, H., Zeng, Y., Zhang, Z., Paterson, N. G., Stansfeld, P. J., Wang, Z., Zhang, Y., Wang, W. & Dong, C. Structural basis of outer membrane protein insertion by the BAM complex. Nature 531, 64–69, doi:10.1038/nature17199 (2016).

12 Bakelar, J., Buchanan, S. K. & Noinaj, N. The structure of the beta-barrel assembly machinery complex. Science (New York, N.Y.) 351, 180–186, doi:10.1126/science.aad3460 (2016).

13 Doerner, P. A. & Sousa, M. C. Extreme Dynamics in the BamA beta-Barrel Seam. Biochemistry 56, 3142–3149, doi:10.1021/acs.biochem.7b00281 (2017).

14 Gessmann, D., Chung, Y. H., Danoff, E. J., Plummer, A. M., Sandlin, C. W., Zaccai, N. R. & Fleming, K. G. Outer membrane beta-barrel protein folding is physically controlled by periplasmic lipid head groups and BamA. Proceedings of the National Academy of Sciences of the United States of America 111, 5878–5883, doi:10.1073/pnas.1322473111 (2014).

15 Doyle, M. T. & Bernstein, H. D. Bacterial outer membrane proteins assemble via asymmetric interactions with the BamA beta-barrel. Nature communications 10, 3358, doi:10.1038/s41467-019-11230-9 (2019).

16 Tomasek, D., Rawson, S., Lee, J., Wzorek, J. S., Harrison, S. C., Li, Z. & Kahne, D. Structure of a nascent membrane protein as it folds on the BAM complex. Nature, doi:10.1038/s41586-020-2370-1 (2020).

17 Sklar, J. G., Wu, T., Gronenberg, L. S., Malinverni, J. C., Kahne, D. & Silhavy, T. J. Lipoprotein SmpA is a component of the YaeT complex that assembles outer membrane proteins in Escherichia coli. Proceedings of the National Academy of Sciences of the United States of America 104, 6400–6405, doi:10.1073/pnas.0701579104 (2007).

18 Kim, S., Malinverni, J. C., Sliz, P., Silhavy, T. J., Harrison, S. C. & Kahne, D. Structure and function of an essential component of the outer membrane protein assembly machine. Science (New York, N.Y.) 317, 961–964, doi:10.1126/science.1143993 (2007).

19 Wu, T., Malinverni, J., Ruiz, N., Kim, S., Silhavy, T. J. & Kahne, D. Identification of a multicomponent complex required for outer membrane biogenesis in Escherichia coli. Cell 121, 235–245, doi:10.1016/j.cell.2005.02.015 (2005).

20 Bennion, D., Charlson, E. S., Coon, E. & Misra, R. Dissection of beta-barrel outer membrane protein assembly pathways through characterizing BamA POTRA 1 mutants of Escherichia coli. Molecular microbiology 77, 1153–1171, doi:10.1111/j.1365-2958.2010.07280.x (2010).

21 Rizzitello, A. E., Harper, J. R. & Silhavy, T. J. Genetic evidence for parallel pathways of chaperone activity in the periplasm of Escherichia coli. Journal of bacteriology 183, 6794–6800, doi:10.1128/jb.183.23.6794-6800.2001 (2001).

22 Crane, J. M. & Randall, L. L. The Sec System: Protein Export in Escherichia coli. EcoSal Plus 7, doi:10.1128/ecosalplus.ESP-0002-2017 (2017).

23 Walsh, N. P., Alba, B. M., Bose, B., Gross, C. A. & Sauer, R. T. OMP peptide signals initiate the envelope-stress response by activating DegS protease via relief of inhibition mediated by its PDZ domain. Cell 113, 61–71, doi:10.1016/s0092-8674(03)00203-4 (2003).

24 Ades, S. E. Regulation by destruction: design of the sigmaE envelope stress response. Current opinion in microbiology 11, 535–540, doi:10.1016/j.mib.2008.10.004 (2008).

25 Rhodius, V. A., Suh, W. C., Nonaka, G., West, J. & Gross, C. A. Conserved and variable functions of the sigmaE stress response in related genomes. PLoS biology 4, e2, doi:10.1371/journal.pbio.0040002 (2006).

26 Guillier, M., Gottesman, S. & Storz, G. Modulating the outer membrane with small RNAs. Genes & development 20, 2338–2348, doi:10.1101/gad.1457506 (2006).

27 De Las Penas, A., Connolly, L. & Gross, C. A. SigmaE is an essential sigma factor in Escherichia coli. Journal of bacteriology 179, 6862–6864, doi:10.1128/jb.179.21.6862-6864.1997 (1997).

28 Missiakas, D., Mayer, M. P., Lemaire, M., Georgopoulos, C. & Raina, S. Modulation of the Escherichia coli sigmaE (RpoE) heat-shock transcription-factor activity by the RseA, RseB and RseC proteins. Molecular microbiology 24, 355–371, doi:10.1046/j.1365-2958.1997.3601713.x (1997).

29 De Las Penas, A., Connolly, L. & Gross, C. A. The sigmaE-mediated response to extracytoplasmic stress in Escherichia coli is transduced by RseA and RseB, two negative regulators of sigmaE. Molecular microbiology 24, 373–385, doi:10.1046/j.1365-2958.1997.3611718.x (1997).

30 Nicoloff, H., Gopalkrishnan, S. & Ades, S. E. Appropriate Regulation of the sigma(E)-Dependent Envelope Stress Response Is Necessary To Maintain Cell Envelope Integrity and Stationary-Phase Survival in Escherichia coli. Journal of bacteriology 199, doi:10.1128/jb.00089-17 (2017).

31 Mitchell, A. M. & Silhavy, T. J. Envelope stress responses: balancing damage repair and toxicity. Nature reviews. Microbiology 17, 417–428, doi:10.1038/s41579-019-0199-0 (2019).

32 Morris, F. C., Wells, T. J., Bryant, J. A., Schager, A. E., Sevastsyanovich, Y. R., Squire, D. J. P., Marshall, J., Isom, G. L., Rooke, J., Maderbocus, R., Knowles, T. J., Overduin, M., Rossiter, A. E., Cunningham, A. F. & Henderson, I. R. YraP Contributes to Cell Envelope Integrity and Virulence of Salmonella enterica Serovar Typhimurium. Infection and immunity 86, doi:10.1128/iai.00829-17 (2018).

33 Tsang, M. J., Yakhnina, A. A. & Bernhardt, T. G. NlpD links cell wall remodeling and outer membrane invagination during cytokinesis in Escherichia coli. PLoS genetics 13, e1006888, doi:10.1371/journal.pgen.1006888 (2017).

34 Onufryk, C., Crouch, M. L., Fang, F. C. & Gross, C. A. Characterization of six lipoproteins in the sigmaE regulon. Journal of bacteriology 187, 4552–4561, doi:10.1128/jb.187.13.4552-4561.2005 (2005).

35 Seib, K. L., Haag, A. F., Oriente, F., Fantappie, L., Borghi, S., Semchenko, E. A., Schulz, B. L., Ferlicca, F., Taddei, A. R., Giuliani, M. M., Pizza, M. & Delany, I. The meningococcal vaccine antigen GNA2091 is an analogue of YraP and plays key roles in outer membrane stability and virulence. FASEB journal : official publication of the Federation of American Societies for Experimental Biology 33, 12324–12335, doi:10.1096/fj.201900669R (2019).

36 Bos, M. P., Grijpstra, J., Tommassen-van Boxtel, R. & Tommassen, J. Involvement of Neisseria meningitidis lipoprotein GNA2091 in the assembly of a subset of outer membrane proteins. The Journal of biological chemistry 289, 15602–15610, doi:10.1074/jbc.M113.539510 (2014).

37 Baba, T., Ara, T., Hasegawa, M., Takai, Y., Okumura, Y., Baba, M., Datsenko, K. A., Tomita, M., Wanner, B. L. & Mori, H. Construction of Escherichia coli K-12 in-frame, single-gene knockout mutants: the Keio collection. Molecular systems biology 2, 2006.0008, doi:10.1038/msb4100050 (2006).

38 Cui, L., Vigouroux, A., Rousset, F., Varet, H., Khanna, V. & Bikard, D. A CRISPRi screen in E. coli reveals sequence-specific toxicity of dCas9. Nature communications 9, 1912, doi:10.1038/s41467-018-04209-5 (2018).

39 Calvo-Villamanan, A., Ng, J. W., Planel, R., Menager, H., Chen, A., Cui, L. & Bikard, D. On-target activity predictions enable improved CRISPR-dCas9 screens in bacteria. Nucleic acids research, doi:10.1093/nar/gkaa294 (2020).

40 Pichoff, S., Du, S. & Lutkenhaus, J. Roles of FtsEX in cell division. Research in microbiology 170, 374–380, doi:10.1016/j.resmic.2019.07.003 (2019).

41 Heidrich, C., Templin, M. F., Ursinus, A., Merdanovic, M., Berger, J., Schwarz, H., de Pedro, M. A. & Holtje, J. V. Involvement of N-acetylmuramyl-L-alanine amidases in cell separation and antibiotic-induced autolysis of Escherichia coli. Molecular microbiology 41, 167–178, doi:10.1046/j.1365-2958.2001.02499.x (2001).

42 Uehara, T., Dinh, T. & Bernhardt, T. G. LytM-domain factors are required for daughter cell separation and rapid ampicillin-induced lysis in Escherichia coli. Journal of bacteriology 191, 5094–5107, doi:10.1128/jb.00505-09 (2009).

43 Yang, D. C., Peters, N. T., Parzych, K. R., Uehara, T., Markovski, M. & Bernhardt, T. G. An ATP-binding cassette transporter-like complex governs cell-wall hydrolysis at the bacterial cytokinetic ring. Proceedings of the National Academy of Sciences of the United States of America 108, E1052–1060, doi:10.1073/pnas.1107780108 (2011).

44 Uehara, T., Parzych, K. R., Dinh, T. & Bernhardt, T. G. Daughter cell separation is controlled by cytokinetic ring-activated cell wall hydrolysis. The EMBO journal 29, 1412–1422, doi:10.1038/emboj.2010.36 (2010).

45 Chung, H. S., Yao, Z., Goehring, N. W., Kishony, R., Beckwith, J. & Kahne, D. Rapid beta-lactam-induced lysis requires successful assembly of the cell division machinery. Proceedings of the National Academy of Sciences of the United States of America 106, 21872–21877, doi:10.1073/pnas.0911674106 (2009).

46 Malinverni, J. C., Werner, J., Kim, S., Sklar, J. G., Kahne, D., Misra, R. & Silhavy, T. J. YfiO stabilizes the YaeT complex and is essential for outer membrane protein assembly in Escherichia coli. Molecular microbiology 61, 151–164, doi:10.1111/j.1365-2958.2006.05211.x (2006).

47 Carlson, M. L., Stacey, R. G., Young, J. W., Wason, I. S., Zhao, Z., Rattray, D. G., Scott, N., Kerr, C. H., Babu, M., Foster, L. J. & Duong Van Hoa, F. Profiling the Escherichia coli membrane protein interactome captured in Peptidisc libraries. eLife 8, doi:10.7554/eLife.46615 (2019).

48 Alvira, S., Watkins, D. W., Troman, L., Allen, W. J., Lorriman, J., Degliesposti, G., Cohen, E. J., Beeby, M., Daum, B., Gold, V. A. M., Skehel, J. M. & Collinson, I. Inter-membrane association of the Sec and BAM translocons for bacterial outer-membrane biogenesis. bioRxiv, 589077, doi:10.1101/589077 (2020).

49 Li, G. W., Burkhardt, D., Gross, C. & Weissman, J. S. Quantifying absolute protein synthesis rates reveals principles underlying allocation of cellular resources. Cell 157, 624–635, doi:10.1016/j.cell.2014.02.033 (2014).

50 Nakamura, K. & Mizushima, S. Effects of heating in dodecyl sulfate solution on the conformation and electrophoretic mobility of isolated major outer membrane proteins from Escherichia coli K-12. Journal of biochemistry 80, 1411–1422, doi:10.1093/oxfordjournals.jbchem.a131414 (1976).

51 Rouviere, P. E. & Gross, C. A. SurA, a periplasmic protein with peptidyl-prolyl isomerase activity, participates in the assembly of outer membrane porins. Genes & development 10, 3170–3182, doi:10.1101/gad.10.24.3170 (1996).

52 Charlson, E. S., Werner, J. N. & Misra, R. Differential effects of yfgL mutation on Escherichia coli outer membrane proteins and lipopolysaccharide. Journal of bacteriology 188, 7186–7194, doi:10.1128/jb.00571-06 (2006).

53 Guo, M. S., Updegrove, T. B., Gogol, E. B., Shabalina, S. A., Gross, C. A. & Storz, G. MicL, a new σE-dependent sRNA, combats envelope stress by repressing synthesis of Lpp, the major outer membrane lipoprotein. Genes & development 28, 1620–1634, doi:10.1101/gad.243485.114 (2014).

54 Gogol, E. B., Rhodius, V. A., Papenfort, K., Vogel, J. & Gross, C. A. Small RNAs endow a transcriptional activator with essential repressor functions for single-tier control of a global stress regulon. Proceedings of the National Academy of Sciences of the United States of America 108, 12875–12880, doi:10.1073/pnas.1109379108 (2011).

55 Roman-Hernandez, G., Peterson, J. H. & Bernstein, H. D. Reconstitution of bacterial autotransporter assembly using purified components. eLife 3, e04234, doi:10.7554/eLife.04234 (2014).

56 Hale, C. A. & de Boer, P. A. Direct binding of FtsZ to ZipA, an essential component of the septal ring structure that mediates cell division in E. coli. Cell 88, 175–185, doi:10.1016/s0092-8674(00)81838-3 (1997).

57 Hartmann, J. B., Zahn, M., Burmann, I. M., Bibow, S. & Hiller, S. Sequence-Specific Solution NMR Assignments of the beta-Barrel Insertase BamA to Monitor Its Conformational Ensemble at the Atomic Level. Journal of the American Chemical Society 140, 11252–11260, doi:10.1021/jacs.8b03220 (2018).

58 Vertommen, D., Ruiz, N., Leverrier, P., Silhavy, T. J. & Collet, J. F. Characterization of the role of the Escherichia coli periplasmic chaperone SurA using differential proteomics. Proteomics 9, 2432–2443, doi:10.1002/pmic.200800794 (2009).

59 Konovalova, A., Schwalm, J. A. & Silhavy, T. J. A Suppressor Mutation That Creates a Faster and More Robust sigmaE Envelope Stress Response. Journal of bacteriology 198, 2345–2351, doi:10.1128/jb.00340-16 (2016).

60 Kern, B., Leiser, O. P. & Misra, R. Suppressor Mutations in degS Overcome the Acute Temperature-Sensitive Phenotype of DeltadegP and DeltadegP Deltatol-pal Mutants of Escherichia coli. Journal of bacteriology 201, doi:10.1128/jb.00742-18 (2019).

61 Hart, E. M., O’Connell, A., Tang, K., Wzorek, J. S., Grabowicz, M., Kahne, D. & Silhavy, T. J. Fine-Tuning of sigma(E) Activation Suppresses Multiple Assembly-Defective Mutations in Escherichia coli. Journal of bacteriology 201, doi:10.1128/jb.00745-18 (2019).

62 Leiser, O. P., Charlson, E. S., Gerken, H. & Misra, R. Reversal of the ΔdegP phenotypes by a novel rpoE allele of Escherichia coli. PloS one 7, e33979, doi:10.1371/journal.pone.0033979 (2012).

63 Yeats, C. & Bateman, A. The BON domain: a putative membrane-binding domain. Trends in biochemical sciences 28, 352–355, doi:10.1016/s0968-0004(03)00115-4 (2003).

64 Bryant, J. A., Morris, F. C., Knowles, T. J., Maderbocus, R., Heinz, E., Boelter, G., Alodaini, D., Colyer, A., Wotherspoon, P. J., Staunton, K. A., Jeeves, M., Browning, D. F., Sevastsyanovich, Y. R., Wells, T. J., Rossiter, A. E., Bavro, V. N., Sridhar, P., Ward, D. G., Chong, Z. S., Icke, C., Teo, A., Chng, S. S., Roper, D. I., Lithgow, T., Cunningham, A. F., Banzhaf, M., Overduin, M. & Henderson, I. R. Structure-function analyses of dual-BON domain protein DolP identifies phospholipid binding as a new mechanism for protein localisation. bioRxiv, 2020.2008.2010.244616, doi:10.1101/2020.08.10.244616 (2020).

65 Fleming, K. G. A combined kinetic push and thermodynamic pull as driving forces for outer membrane protein sorting and folding in bacteria. Philosophical transactions of the Royal Society of London. Series B, Biological sciences 370, doi:10.1098/rstb.2015.0026 (2015).

66 Horne, J. E., Brockwell, D. J. & Radford, S. E. Role of the lipid bilayer in outer membrane protein folding in Gram-negative bacteria. The Journal of biological chemistry 295, 10340–10367, doi:10.1074/jbc.REV120.011473 (2020).

67 Sinnige, T., Weingarth, M., Renault, M., Baker, L., Tommassen, J. & Baldus, M. Solid-state NMR studies of full-length BamA in lipid bilayers suggest limited overall POTRA mobility. Journal of molecular biology 426, 2009–2021, doi:10.1016/j.jmb.2014.02.007 (2014).

68 Rassam, P., Copeland, N. A., Birkholz, O., Tóth, C., Chavent, M., Duncan, A. L., Cross, S. J., Housden, N. G., Kaminska, R., Seger, U., Quinn, D. M., Garrod, T. J., Sansom, M. S., Piehler, J., Baumann, C. G. & Kleanthous, C. Supramolecular assemblies underpin turnover of outer membrane proteins in bacteria. Nature 523, 333–336, doi:10.1038/nature14461 (2015).

69 Gunasinghe, S. D., Shiota, T., Stubenrauch, C. J., Schulze, K. E., Webb, C. T., Fulcher, A. J., Dunstan, R. A., Hay, I. D., Naderer, T., Whelan, D. R., Bell, T. D. M., Elgass, K. D., Strugnell, R. A. & Lithgow, T. The WD40 Protein BamB Mediates Coupling of BAM Complexes into Assembly Precincts in the Bacterial Outer Membrane. Cell reports 23, 2782–2794, doi:10.1016/j.celrep.2018.04.093 (2018).

70 Jarosławski, S., Duquesne, K., Sturgis, J. N. & Scheuring, S. High-resolution architecture of the outer membrane of the Gram-negative bacteria Roseobacter denitrificans. Molecular microbiology 74, 1211–1222, doi:10.1111/j.1365-2958.2009.06926.x (2009).

71 Lessen, H. J., Fleming, P. J., Fleming, K. G. & Sodt, A. J. Building Blocks of the Outer Membrane: Calculating a General Elastic Energy Model for β-Barrel Membrane Proteins. Journal of chemical theory and computation 14, 4487–4497, doi:10.1021/acs.jctc.8b00377 (2018).

72 Cascales, E., Bernadac, A., Gavioli, M., Lazzaroni, J. C. & Lloubes, R. Pal lipoprotein of Escherichia coli plays a major role in outer membrane integrity. Journal of bacteriology 184, 754–759, doi:10.1128/jb.184.3.754-759.2002 (2002).

73 Asmar, A. T. & Collet, J. F. Lpp, the Braun lipoprotein, turns 50-major achievements and remaining issues. FEMS microbiology letters 365, doi:10.1093/femsle/fny199 (2018).

74 Grenier, F., Matteau, D., Baby, V. & Rodrigue, S. Complete Genome Sequence of Escherichia coli BW25113. Genome announcements 2, doi:10.1128/genomeA.01038-14 (2014).

75 Blattner, F. R., Plunkett, G., 3rd, Bloch, C. A., Perna, N. T., Burland, V., Riley, M., Collado-Vides, J., Glasner, J. D., Rode, C. K., Mayhew, G. F., Gregor, J., Davis, N. W., Kirkpatrick, H. A., Goeden, M. A., Rose, D. J., Mau, B. & Shao, Y. The complete genome sequence of Escherichia coli K-12. Science (New York, N.Y.) 277, 1453–1462, doi:10.1126/science.277.5331.1453 (1997).

76 Datsenko, K. A. & Wanner, B. L. One-step inactivation of chromosomal genes in Escherichia coli K-12 using PCR products. Proceedings of the National Academy of Sciences of the United States of America 97, 6640–6645, doi:10.1073/pnas.120163297 (2000).

77 Knop, M., Siegers, K., Pereira, G., Zachariae, W., Winsor, B., Nasmyth, K. & Schiebel, E. Epitope tagging of yeast genes using a PCR-based strategy: more tags and improved practical routines. Yeast (Chichester, England) 15, 963–972, doi:10.1002/(sici)1097-0061(199907)15:10b<963::Aid-yea399>3.0.Co;2-w (1999).

78 McLeay, R. C. & Bailey, T. L. Motif Enrichment Analysis: a unified framework and an evaluation on ChIP data. BMC bioinformatics 11, 165, doi:10.1186/1471-2105-11-165 (2010).

79 Ieva, R. & Bernstein, H. D. Interaction of an autotransporter passenger domain with BamA during its translocation across the bacterial outer membrane. Proceedings of the National Academy of Sciences of the United States of America 106, 19120–19125, doi:10.1073/pnas.0907912106 (2009).

80 Vischer, N. O., Verheul, J., Postma, M., van den Berg van Saparoea, B., Galli, E., Natale, P., Gerdes, K., Luirink, J., Vollmer, W., Vicente, M. & den Blaauwen, T. Cell age dependent concentration of Escherichia coli divisome proteins analyzed with ImageJ and ObjectJ. Frontiers in microbiology 6, 586, doi:10.3389/fmicb.2015.00586 (2015).

81 Ashburner, M., Ball, C. A., Blake, J. A., Botstein, D., Butler, H., Cherry, J. M., Davis, A. P., Dolinski, K., Dwight, S. S., Eppig, J. T., Harris, M. A., Hill, D. P., Issel-Tarver, L., Kasarskis, A., Lewis, S., Matese, J. C., Richardson, J. E., Ringwald, M., Rubin, G. M. & Sherlock, G. Gene ontology: tool for the unification of biology. The Gene Ontology Consortium. Nature genetics 25, 25–29, doi:10.1038/75556 (2000).

82 Albenne, C. & Ieva, R. Job contenders: roles of the beta-barrel assembly machinery and the translocation and assembly module in autotransporter secretion. Molecular microbiology 106, 505–517, doi:10.1111/mmi.13832 (2017).

83 Szabady, R. L., Peterson, J. H., Skillman, K. M. & Bernstein, H. D. An unusual signal peptide facilitates late steps in the biogenesis of a bacterial autotransporter. Proceedings of the National Academy of Sciences of the United States of America 102, 221–226, doi:10.1073/pnas.0406055102 (2005).

